# Ral signals through a MAP4 Kinase-p38 MAP kinase cascade in *C. elegans* cell fate patterning

**DOI:** 10.1101/381707

**Authors:** Hanna Shin, Rebecca E.W. Kaplan, Tam Duong, Razan Fakieh, David J. Reiner

**Affiliations:** Center for Translational Cancer Research, Institute of Biosciences and Technology, Texas A&M Health Science Center, Houston, TX, 77030, USA.; Department of Medical Physiology, College of Medicine, Texas A&M University, College Station, TX, 77843, USA.; Department of Pharmacology and Lineberger Comprehensive Cancer Center, University of North Carolina, Chapel Hill, NC, 27599, USA.

**Keywords:** MAP4K, Sec5, RalBPI, RLBP-1, EXOC-8, SEC-5, MLK-1, MIG-15

## Abstract

*C. elegans* vulval precursor cell (VPC) fates are patterned by an EGF gradient. High dose EGF induces 1° VPC fate, while lower dose EGF contributes to 2° fate in support of LIN-12/Notch. We previously showed that the EGF 2°-promoting signal is mediated by LET-60/Ras switching effectors, from the canonical Raf-MEK-ERK MAP kinase cascade that promotes 1° fate to the non-canonical RalGEF-Ral that promotes 2° fate. Of oncogenic Ras effectors, RalGEF-Ral is by far the least well-understood. We use genetic analysis to identify an effector cascade downstream of *C. elegans* RAL-1/Ral, starting with an established Ral binding partner, Exo84 of the exocyst complex. Additionally, RAL-1 signals through GCK-2, a CNH domain-containing MAP4 kinase, and PMK-1/p38 MAP kinase cascade to promote 2° fate. Our study delineates a Ral-dependent developmental signaling cascade *in vivo*, thus providing the mechanism by which lower EGF dose is transduced.

## Introduction

Ras is the most mutated oncoprotein. Yet strategies to inhibit oncogenic Ras have failed, so Ras is considered to be mostly “undruggable” (Papke and Der, 2017). Consequently, attention has shifted to oncogenic Ras effectors to identify therapeutic targets. Canonical oncogenic Ras effectors, the Raf-MEK-ERK and PI3K-PDK-Akt cascades, are among the best studied and most targeted signaling cascades (Ryan et al., 2015; Wong et al., 2010). Yet even potent small molecule inhibitors, like the BRAF inhibitor vemurafenib, are subject to multiple bypass mechanisms that permit initially responsive tumors to relapse (Sun et al., 2014). Thus, successful treatment will likely require multi-pronged regimens to simultaneously inhibit multiple Ras effectors.

In addition to the canonical Raf and PI3K cascades, Ras uses RalGEF-Ral to promote tumorigenesis (Feig, 2003). Historically, canonical Ras-Raf and Ras-PI3K signaling was shown to cause cancer transformation of mouse primary fibroblasts (Khosravi-Far et al., 1996; Kyriakis et al., 1992; White et al., 1995). The emergence of immortalized human epithelial cell culture led to the key finding that Ras-RalGEF-Ral is also a critical player in human oncogenesis (Hamad et al., 2002; Urano et al., 1996; White et al., 1996). RalGEF is an exchange factor that promotes GTP loading of the Ral (<u>Ra</u>s <u>l</u>ike) small GTPase (Feig, 2003). Loss of RalGAP (Ral <u>G</u>TPase <u>a</u>ctivating <u>p</u>rotein), a putative tumor suppressor, increases tumorigenesis without activated Ras (Oeckinghaus et al., 2014; Saito et al., 2013), further supporting the importance of Ral signaling in cancer.

Three binding partners of Ral have been well validated: RalBP1 (Ral binding protein 1) and Sec5 and Exo84 subunits of the heterooctameric exocyst complex (reviewed in Gentry et al., 2014; Fig. S1). The exocyst represents an unusual roadblock to biochemical bootstrapping of signaling activities: the exocyst is broadly integral to essential cell biological processes (e.g. exocytosis, PAR/polarity complex; Wu and Guo, 2015) and potentially binds to hundreds of partners, thus mostly precluding identification of downstream signaling partners via binding studies. Consequently, beyond these immediate binding partners, we know little of downstream functions of Ral signaling through the exocyst *in vivo*.

Studies in *Drosophila* provided key hints to the nature of Ral downstream signaling in development. In morphogenetic events, DRal was implicated in antagonizing the JNK MAP kinase (Sawamoto et al., 1999). In bristle apoptosis assays, DRal was found to have negative and positive relationships with JNK and p38 MAP kinase cascades, respectively (Balakireva et al., 2006). Importantly, this study also established that exocyst component Sec5 binds to HGK/NIK/MAPK4, a CNH domain containing MAP4 kinase. The *Drosophila* ortholog, Msn (Misshapen) was found to function antagonistically to DRal (Balakireva et al., 2006), and is known to function with JNK in *Drosophila* embryonic dorsal closure and other morphogenetic events (Su et al., 1998). Yet these studies relied on ectopic over-expression and dominant-negative reagents, which complicated interpretation. Furthermore, direct genetic epistasis could not be assayed because many of the proteins studied are essential for development in *Drosophila*.

The Ste20 family of mitogen-activated protein kinase kinase kinase kinases (MAP4 Kinases or MAP4Ks) is conserved throughout eukaryotes (Dan et al., 2001; Delpire, 2009). Two paralogous subfamilies of this group, GCK-I and GCK-IV (<u>G</u>erminal <u>C</u>enter <u>K</u>inases), are defined by distinctive domain architecture: an N-terminal S/T kinase domain, a C-terminal CNH domain (for <u>C</u>itron <u>N</u>-terminal <u>H</u>omology), and an unstructured poly-proline linker region (Fig. 1A; Dan et al., 2001). *C. elegans* GCK-2 (ZC404.9) is an 829-residue protein in the GCK-I subfamily (the “GCK-2 group”: *Drosophila* Hppy (Happyhour), mammalian MAP4K1/HPK1, MAP4K2/GCK, MAP4K3/GLK, and MAP4K5/GCKR/KHS1; Figs. S1B, S1C). *C. elegans* MIG-15 is in the GCK-IV subfamily (the “MIG-15 group”: *C. elegans* MIG-15, *Drosophila* Msn, mammalian MAP4K4/NIK/HGK, MAP4K6/MINK, MAP4K7/TNIK, and MAP4K8/NRK/NESK; Figs. S1D, S1E).

**Figure 1.**
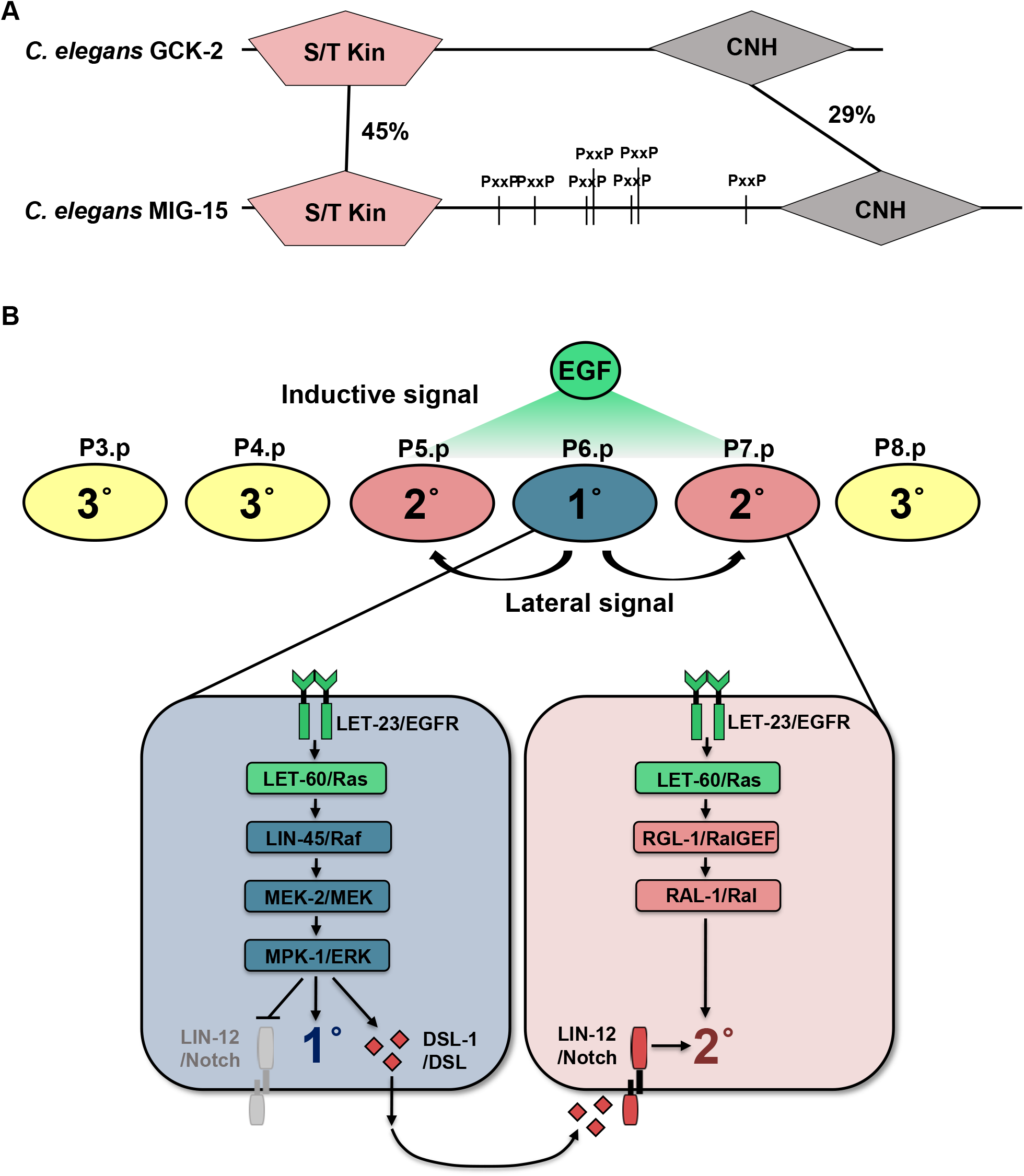
*C. elegans* CNH-domain organization MAP4Ks and VPC fate patterning. Domain organization and conservation of paralogous *C. elegans* MAP4Ks GCK-2 and MIG-15. (B) VPCs, P3.p through P8.p, are patterned to assume 3°-3°-2°-1°-2°-3° fate by coordinated graded action of EGF secreted from the anchor cell and Notch lateral signal. LET-60/Ras-LIN-45/Raf-MEK-2/MEK-MPK-1/ERK and LET-60/Ras-RGL-1/RalGEF-RAL-1/Ral promote 1° fate and 2° induction, respectively. Presumptive 1° cells synthesize DSL ligands to induce 2° fate via LIN-12/Notch.

**Figure 2.**
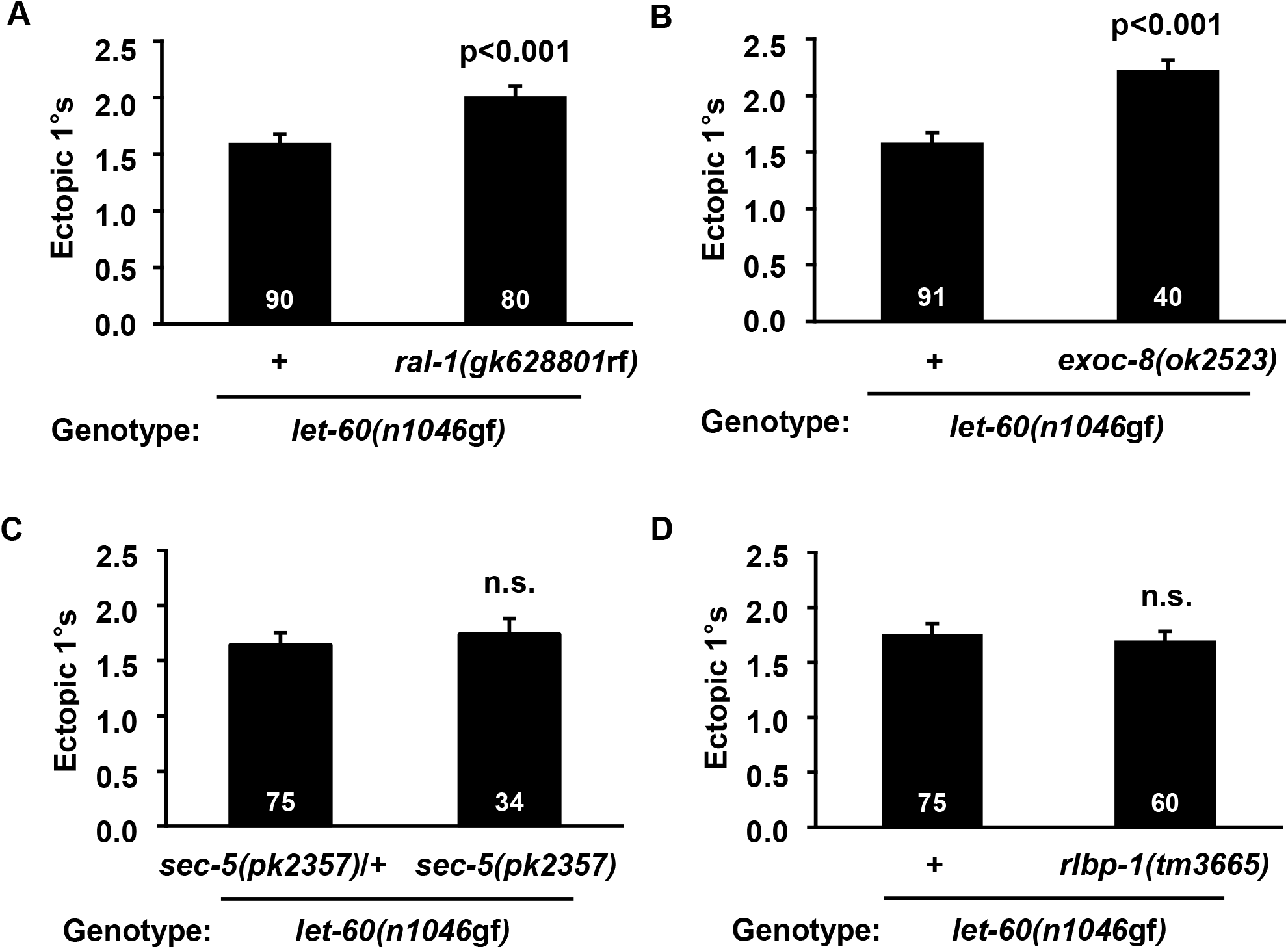
EXOC-8 functions in VPC fate patterning. The Y-axis indicates number of ectopic 1° cells. The (A) *ral-1 (gk628801*rf) R139H missense and (B) *exoc-8(ok2523)* deletion mutations increased ectopic 1° induction in the *let-60(n1046*gf) background. (C) The sec-*5(pk2358)* late nonsense allele (Frische et al. 2007) did not alter ectopic 1 ° induction in the *let-60(n1046*gf) background. (D) The *rlbp-1* out-of-frame deletion allele, *tm3665*, conferred no change in the *let-60(n1046*gf) background. N indicated in white on columns. P value calculated by *t* test. Error bars = S.E.M.

Critically, *Drosophila* Hppy was as yet undiscovered at the time of Msn investigation relative to DRal (Balakireva et al., 2006). Hppy antagonizes canonical EGFR signaling through ERK MAPK in ethanol response; its relationship to DRal was not studied (Corl et al., 2009). Thus, we turned to *C. elegans* VPC fate patterning to investigate a signaling cascade downstream of Ral relative to these enigmatic MAP4 kinases.

During the L3 stage, EGF produced by the gonadal Anchor Cell (AC) induces six initially equipotent Vulval Precursor Cells (VPCs), P3.p through P8.p, to assume the highly reproducible 3°-3°-2°-1°-2°-3° pattern (Fig. 1B). 1° and 2° cells undergo stereotyped divisions and morphogenesis to form the mature vulva, while uninduced 3° cells divide once and fuse with surrounding cells (Sternberg, 2005). Historically, two competing models, the “Morphogen Gradient Model” and the “Sequential Induction Model”, were posited to describe VPC fate patterning. In the “Morphogen Gradient Model,” graded inductive signal controls fate patterning: the VPC closest to the AC (typically P6.p) receives the highest LIN-3/EGF-LET-23/EGFR signal to induce 1 ° fate, while neighboring VPCs, P5.p and P7.p, receive lower LIN-3/EGF-LET-23/EGFR signal, and thus become 2° (Katz et al., 1995; Katz et al., 1996; Sternberg and Horvitz, 1986, 1989). Yet identification of key genes in VPC patterning led to the potentially contradictory “Sequential Induction Model.” LET-23/EGFR and LIN-12/Notch are necessary and sufficient for 1° and 2° induction, respectively (Aroian et al., 1990; Greenwald et al., 1983). Activation of LET-23/EGFR triggers a LIN-45/Raf-MEK-2/MEK-MPK-1/ERK canonical MAP kinase (MAPK) cascade to induce 1° fate (reviewed in Sundaram, 2013). These presumptive 1 ° cells in turn secrete DSL ligands to induce neighbors to become 2° via the LIN-12/Notch (Chen and Greenwald, 2004). LET-23/EGFR was found to function cell autonomously to induce 1 ° fate, further supporting the “Sequential Induction Model” (Koga and Ohshima, 1995; Simske and Kim, 1995). The two models long remained unreconciled, and no mechanism was known by which graded LIN-3/EGF-LET-23/EGFR activity promotes 2° fate (Kenyon, 1995).

Overlaid on this system are “Mutual Antagonism” mechanisms by which, after initial induction, presumptive 1° and 2° cells enact programs to exclude potentially contradictory signals. For example, in presumptive 1° cells, LIN-12/Notch receptor is internalized and degraded to prohibit conflicting 2°-promoting signaling (Shaye and Greenwald, 2002, 2005). Conversely, in presumptive 2° cells LIN-12/Notch-dependent transcription of LIP-1/ERK phosphatase impedes conflicting 1 °-promoting MPK-1/ERK signaling (Berset et al., 2001; Yoo et al., 2004). Such antagonistic signals are proposed to act collectively to transition from initial patterning specification to commitment (Sternberg, 2005), thereby avoiding inappropriate and/or ambiguous cell fates that can result from inappropriate signals.

Given the importance of Ras-RalGEF-Ral signaling in cancer, we set out to define a role for this signaling module in VPC fate patterning. We found that LET-60/Ras uses the non-canonical RGL-1/RalGEF-RAL-1/Ral effector to promote 2° fate in support of LIN-12/Notch (Zand et al., 2011). Thus, both the Sequential Induction and Morphogen Gradient models are correct: LET-60/Ras switches effectors to interpret the EGF gradient. This mechanism reconciled the two competing models and established a platform for the *in vivo* study of RAL-1/Ral signaling in VPC fate patterning (Reiner, 2011; Zand et al., 2011) (Fig. 1B). Yet the downstream output of the LET-60/Ras-RGL-1/RalGEF-RAL-1/Ral 2°-promoting signal remained unknown.

In this study, we determine that EXOC-8/Exo84, a well-validated Ral-binding protein in mammalian cells, is required to propagate the RAL-1 2°-promoting signal. Significantly, we find that RAL-1 requires the CNH domain-containing GCK-2/MAP4K and PMK-1/p38 MAPK to promote 2° fate. Genetic perturbation of components of this cascade phenocopied perturbation of RAL-1, and these components are necessary for the 2°-promoting activity of mutationally activated RAL-1. 2°-promoting EGF signal requires GCK-2, putative mutationally activated endogenous GCK-2 is sufficient to increase ectopic 2° cell induction, and GCK-2 functions cell autonomously in the VPCs. Using CRISPR/Cas9-dependent genome engineering to tag endogenous gene products with fluorescent protein (FP) and epitope, we observed expression and subcellular localization of endogenous RAL-1, GCK-2, and PMK-1 proteins in VPCs. Our *in vivo* analysis connects Ral to a novel effector cascade in *C. elegans* VPC fate patterning.

## Results

### Criteria for a RAL-1-dependent 2-promoting signal

Since LIN-12/Notch, but not the LET-60/Ras-RGL-1/RalGEF-RAL-1/Ral signal, is necessary for 2° fate induction, we used a combination of parallelism and epistasis to test the genetic relationships among members of the VPC fate-patterning network. Specifically, for this genetic analysis we used two sensitized genetic backgrounds (Fig. S1F). The *let-60(n1046*gf) G13E activating mutation confers excess 1° induction, levels of which are sensitive to perturbation of both 1°- and 2°-promoting signals. The weakly activating *lin-12(n379d)/Notch* mutation both causes ectopic 2° cells and abrogates development of the AC. Consequently, *lin-12(n379*d) provides a simplified signaling milieu in which EGF is not present (Greenwald et al., 1983). This background is sensitive to perturbation of 2°-but not 1°-promoting signals and responds to the EGF-dependent 2°-promoting signal (Zand et al., 2011).

Partly by using these tools, we developed a set of expectations for RAL-1 2°-promoting effectors. (1) Loss of effector function should phenocopy loss of *ral-1* function. (2) Constitutively activated effector should phenocopy constitutively activated RAL-1. (3) Loss of effector function should be epistatic to constitutively activated RAL-1. (4) The effector should function cell autonomously in the VPCs. (5) The effector should be expressed in the VPCs. Using these criteria, we systematically evaluated a putative RAL-1 signaling cascade in 2° VPC fate induction.

### EXOC-8 functions in VPC fate patterning

We tested whether known Ral binding partners, Sec5 and Exo84 of the exocyst complex and RalBP1/RLIP76 (reviewed in Gentry et al., 2014; Fig. S1A), met our first criterion for a RAL-1 effector: loss of effector function should phenocopy loss of *ral-1* function. The heterooctameric exocyst complex generally consists of eight subunits used in different contexts: Sec3, Sec5, Sec6, Sec8, Sec10, Sec15, Exo70 and Exo84 (Wu and Guo, 2015), all of which have single conserved orthologs in *C. elegans*.

Deletion of *ral-1* leads to defects in cell polarity, apparently by disruption of the exocyst complex (Armenti et al., 2014), consistent with previous observations that mammalian Ral functions as a membrane-tethering member of the exocyst (Issaq et al., 2010; Moskalenko et al., 2002; Moskalenko et al., 2003). We previously found that *ral-1(RNAi)* alone did not confer significant vulval patterning defects, nor did a non-null intronic deletion allele of *ral-1* that conferred sterility (Zand et al., 2011). We characterized the *ral-1(gk628801*rf) R139H mutation, which did not confer visible defects (Shin *et al*., in preparation). In the *let-60(n1046*gf) background, *gk628801rf* caused increased 1° induction, consistent with a reduced function (rf) but not null allele of *ral-1*. This result validated our previous findings that reduced *ral-1* signaling and hence reduced 2° signaling increased 1°-promoting signals (Figs. 2A, S2A).

In the same genetic background, we tested effects of the *sec-5(pk2357)* strong hypomorph (Frische et al., 2007), *exoc-8(ok2523)*, and *rlbp-1(tm3665)* deletion alleles *(exoc-7(ok2006)* was included as a negative exocyst control). Among these candidate RAL-1 effectors, only *exoc-8(ok2523)*, a deletion allele, phenocopied *ral-1(gk628801*rf) (Figs. 2B-D, S2B). As expected, *ral-1(gk628801*rf) caused no phenotypic changes in the *lin-12(n379*d) background (Fig. S2C). These results are consistent with EXOC-8 mediating RAL-1 2°-promoting signal.

We also tested *exoc-8(ok2523)* in the *lin-12(n379*d) background. We observed increased ectopic 2° induction (Fig. S2D). This result is inconsistent with our expectation of a RAL-1 effector. Thus, EXOC-8 performs multiple functions in VPC fate patterning, one of which could include mediating a RAL-1 2°-promoting signal.

### Loss of GCK-2 but not MIG-15 confers the same phenotype as loss of RAL-1

To further explore our first criterion, we used *C. elegans* genetics to determine relationships among Ral and the paralogous “GCK-2 Group” and “MIG-15 Group” MAP4Ks. *gck-2(ok2867)* is an in-frame 549 bp deletion in exon 5, resulting in deletion of a part of the kinase domain. *gck-2(tm2537)* is an out-of-frame 565 bp deletion in exon 5, resulting in early stop codons (Khan et al., 2012). Most genetics were done using these alleles and *gck-2-* directed bacterially mediated RNAi (clone V-4P08; Fig. 3A). *mig-15(rh148)* is a V168E missense mutation predicted to abolish kinase function. *mig-15(rh80)* is a W898* late nonsense mutation thought to confer strong loss of function (Chapman et al., 2008). We also used mig-15-directed bacterially mediated RNAi (clone X-5G21).

**Figure 3.**
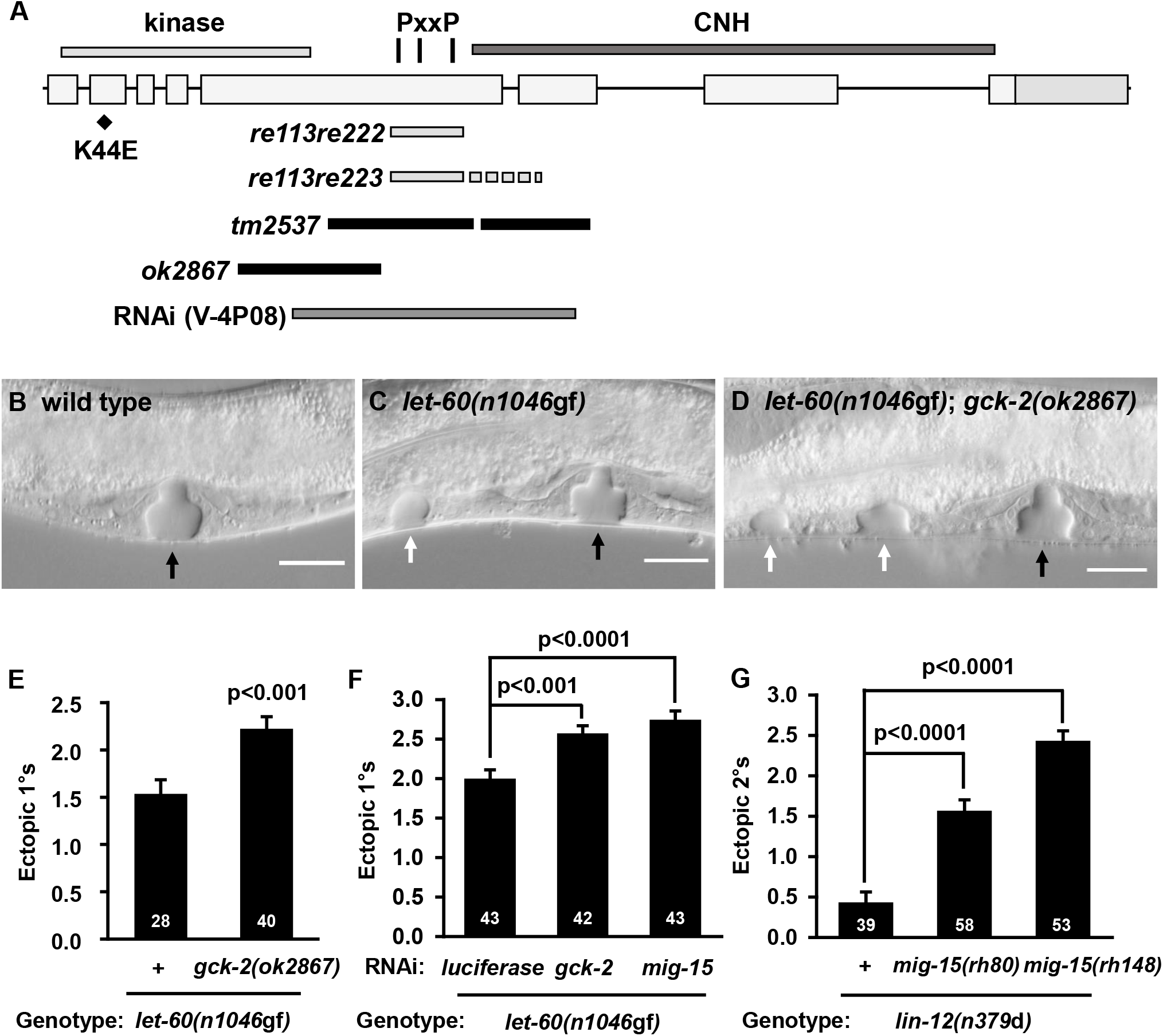
Loss of GCK-2 but not MIG-15 confers defects consistent with a RAL-1 effector. (A) *gck-2* gene structure, domain location, and genetic tools. Light gray: the *re113re222* (312 bp deletion) in-frame and *re113re223* (312 bp deletion, 35 bp insertion) out-of-frame mid-deletions remove the three PxxP sites. Black: *tm2537* (565 bp out-of-frame deletion) and *ok2867* (549 bp in-frame deletion). Dark gray: RNAi target sequence (Kamath et al. 2003). (BD) DIC images of late L4 vulvae and ectopic pseudovulvae. (B) N2 wild type (C) *let-60(n1046*gf), (D) *let-60(n1046gf); gck-2(ok2867)*. Black arrow indicates normal vulva, white arrow indicates ectopic 1° pseudovulvae. Scale bar = 20 μm. (E) *gck-2(ok2867)* enhances ectopic 1° induction in the *let-60(n1046*gf) background. (F) *gck-2(RNAi)* (V-4P08) and *mig-15(RNAi)* (X-5G21) both increase 1° induction in the *let-60(n1046*gf) background. (G) *mig-15(rh80)* and *mig-15(rh148)* do alter ectopic 2° induction in the *lin-12(n379*d) background. P values calculated by *t* test (E) or ANOVA (F, G). Error bars = S.E.M.

*gck-2(ok2867)* conferred increased 1° induction in the *let-60(n1046*gf) background (Figs. 3B-E), similar to *ral-1(gk628801*rf) or *exoc-8(ok2523). gck-2(RNAi)* conferred a phenotype similar to that of *ral-1(RNAi)* (Fig. 3F vs. Fig. S2A). Also, *mig-15(RNAi)* conferred increased 1° induction in the *let-60(n1046*gf) background (Fig. 3F). We also tested the *mig-15(rh148)* reduced function allele in the *let-60(n1046*gf) background. However, we observed severe vulval morphogenesis defects: 2° cells/lineages failed to migrate to join the 1 ° cell/lineage (Fig. S3A). Thus, in double mutant animals we could not discriminate between ectopic 1°s and 2°s that had failed to join the 1° of the normal vulva, which precluded interpretation of VPC induction in these strains (Fig. S3B). Taken together, these results are consistent with both GCK-2 and MIG-15 functioning as either 2°-promoting or 1°-antagonizing signals.

We assessed the roles of MIG-15 and GCK-2 in the *lin-12(n379*d) background (AC/EGF absent, mild ectopic 2° induction). Neither *ral-1(gk628801*rf) nor *gck-2(ok2867* or *tm2537)* altered 2° induction in the *lin-12(n379*d) background (Figs. S2C, 3SC, S3D, respectively). In marked contrast, reduction of *mig-15* function robustly elevated 2° induction in the *lin-12(n379*d) background (Fig. 3G, S3E; the strongest *mig-15* allele, *rh326*, was not assayed due to poor viability). This assay was possible with *mig-15* alleles because the isolated 2°s in the *n379* background could be readily identified, unlike as in the *n1046* background, above.

To test background specificity, we evaluated the impact of *gck-2(ok2867)* in the *lin-3/EGF(n378*rf) hypo-induced rather than the *let-60(n1046*gf) hyper-induced background. *lin-3(n378*rf) supports 20% vulval induction (Hill and Sternberg, 1992). *gck-2(ok2867)* conferred increased vulval induction (Fig. S3F), suggesting that the 2°-promoting signal of GCK-2 is not *let-60(n1046*gf) background dependent.

Collectively, these genetic results are consistent with the hypothesis that GCK-2 functions as an effector of RAL-1 2°-promoting activity. Meeting our first criterion, GCK-2 antagonizes 1° signal and is neutral in the absence of 2°-promoting EGF. This interpretation is supported below by further genetic analysis.

Conversely, MIG-15 is not consistent with a simple criterion of a RAL-1 2°-promoting effector. Unlike RAL-1, MIG-15 antagonizes both 1° and 2° signals, even in the absence of EGF. We cannot exclude MIG-15 as a second RAL-1 effector, perhaps in a negative regulatory relationship, which we will address in the Discussion. Thus, further analysis of MIG-15 function is outside the scope of this study.

### GCK-2 is sufficient to induce 2° fate in support of LIN-12/Notch

Our second criterion is that activated effector should phenocopy constitutively activated RAL-1. RAL-1 is sufficient to promote 2° fate: transgenic VPC-expressed *ral-1(*gf) significantly increased ectopic 2° induction in the *lin-12(n379*d) background (Zand et al., 2011). This same extrachromosomal array, when integrated as *rels10[ral-1(gf)]* (Shin *et al*., in preparation), also increased ectopic 2°s in the *lin-12(n379*d) background (Fig. 4A). Using CRISPR/Cas9-mediated genome editing, we generated a gain-of-function mutation (G26V) in the endogenous *ral-1* locus, also including an in-frame 5’-end mKate2^3xFlag tag (see Figs. S6I, S6J). The resulting *ral-1(re160gf[mKate2^3xFlag::ral-1(G26V)])* caused increased 2° fate induction in the *lin-12(n379*d) background (Figs. 4B-D).

**Figure 4.**
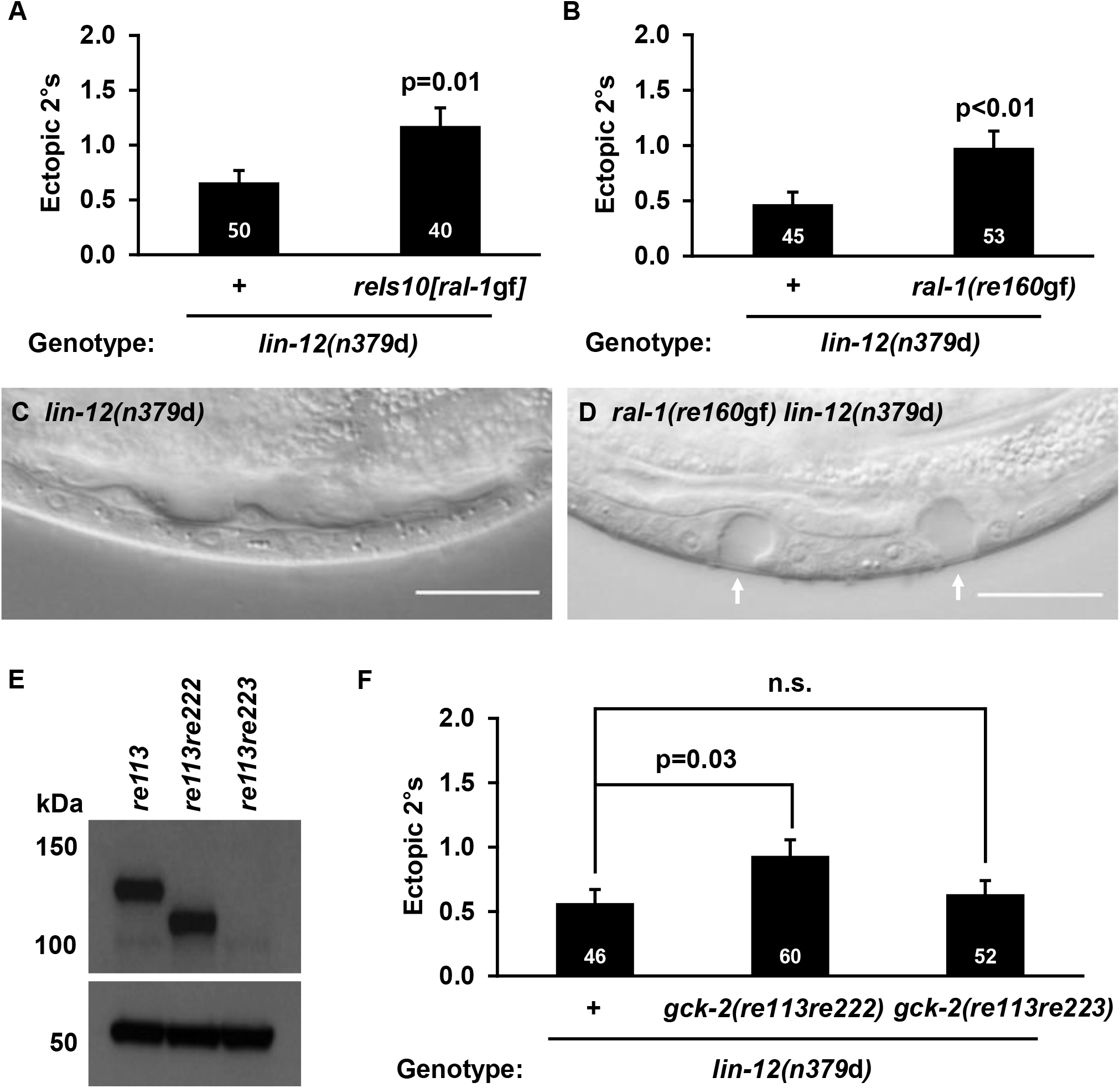
RAL-1 and GCK-2 are sufficient to drive 2° fate induction. (A) Exogenous *(rels10[P_lin-31_::ral-1(gf)])* and (B) endogenous *(ral-1 (re160gf[mKate2^3xFlag::ral-1(G26V)]))* activated RAL-1 increase 2° induction in the *lin-12(n379*d) background. (A, B) Ectopic 2° induction is on the Y axis. (C) Vulvaless (Vul) late L4 *lin-12(n379*d) animal. (D) Ectopic 2° pseudovulvae (white arrows) in *ral-1(re160gf) lin-12(n379*d). Scale bar = 20 μm. (E) Western blot detection of GCK-2 from lysates from endogenously tagged wild-type *(re113)*, in-frame mid-deletion *(re113re222)*, and out-of-frame mid-deletion *(re113re223)* animals, detected by anti-Flag antibody (1:2000). The mNG::3xFlag::GCK-2 fusion protein is predicted to be ~124 kDa, the mid deletion ~110 kDa. (F) The in-frame mid-deletion *gck-2(re113re222)* but not the out-of-frame mid-deletion *gck-2(re113re223)* caused increased 2° induction in the *lin-12(n379*d) background. Ectopic 2° induction is on the Y axis. P value calculated by *t* test or ANOVA. Error bars = S.E.M.

To test whether GCK-2 is sufficient to induce 2° fate, we used CRISPR to generate a putative *gck-2(*gf). Deletion of the proline-rich linker is thought to constitutively activate *Drosophila* Msn (“MIG-15 group”; Su et al., 2000). We deleted the linker in GCK-2 by using the co-CRISPR strategy (Arribere et al., 2014). The *gck-2(re113re222)* mid-Δ was generated in *gck-2(re113[mNG^3xFlag::gck-2])*, which we had already engineered (see below and Figs. 4E, S4A). We also generated the *gck-2(re113re223)* out-of-frame mid-Δ, which served as a negative control. We assessed protein expression in these alleles by western blot: the out-of-frame *re113re223* but not the in-frame *re113re222* abolished detectable tagged GCK-2 (Fig. 4E). The in-frame *re113re222* but not the out-of-frame *re113re223* significantly increased ectopic 2° induction in the *lin-12(n379*d) background (Fig. 4F).

We also tested GCK-2 cell autonomy by generating transgenes expressing VPC-specific putative activating gck-2(mid-Δ) into the *lin-12(n379*d) background, where we observed weakly increased ectopic 2° induction (p = 0.06; Fig. S4B). The same transgene in the *lin-12(n379d); gck-2(tm2537)* background significantly increased ectopic 2° induction (p = 0.03; Fig. S4C). Thus, GCK-2 is sufficient to induce increased 2° induction in support of LIN-12/Notch, consistent with GCK-2 functioning as a 2°-promoting effector of RAL-1.

### GCK-2 functions downstream of LIN-3/EGF and RAL-1

Our third criterion is that loss of effector function should be epistatic to constitutively activated RAL-1. *rels10[ral-1(gf)]* enhanced ectopic 2° induction in the *lin-12(n379*d) background (Fig. 4A) and was blocked by *gck-2(ok2867)* and *gck-2(tm2537)* (Fig. 5A). Thus, GCK-2 meets our third criterion for a RAL-1 effector.

**Figure 5.**
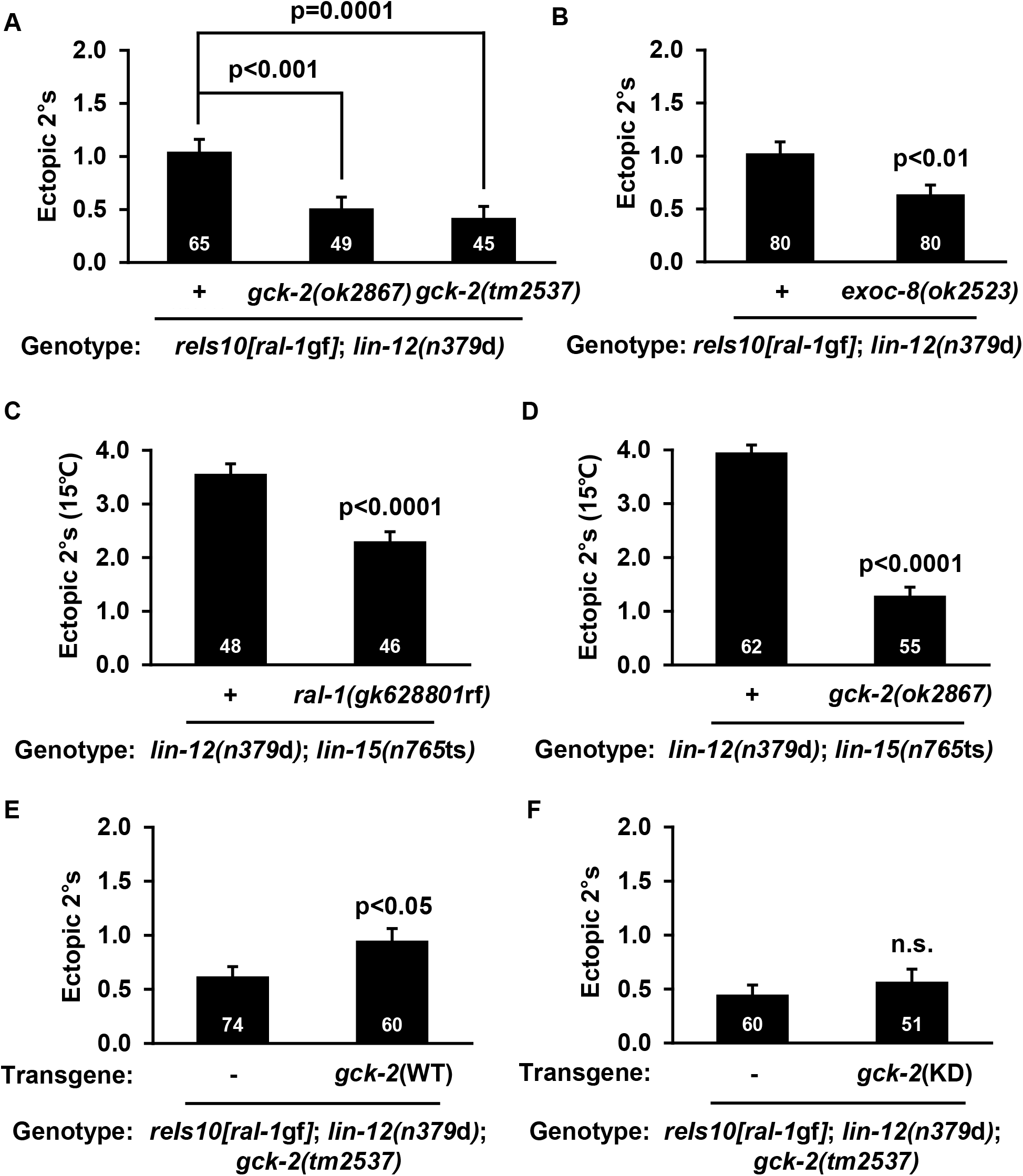
GCK-2 functions cell autonomously downstream of Ral. (A) *gck-2(ok2867)* and *gck-2(tm2537)* blocked the 2°-promoting activity of *rels10[P_lin-31_::ral-1(gf)]* in the *lin-12(n379*d) background, as does (B) *exoc-8(ok2523)*. (C) Strong enhancement of *lin-12(n379d)-* dependent 2° induction by *lin-15(n765ts)* at 15° is reduced by *ral-1(gk628801*rf), and (D) *gck-2(ok2867)*. (E) Vulva-specific expression of wild-type *(reEx176)* but not (F) K44E putative kinase dead *(reEx181)* GCK-2 rescues the suppression of *rels10[ral-1(gf)]* by *gck-2(tm2537)* in the *lin-12(n379*d) background. Each column pair compares array-bearing vs. non-array-bearing siblings. P value calculated by *t* test or ANOVA. Error bars = S.E.M.

**Figure 6.**
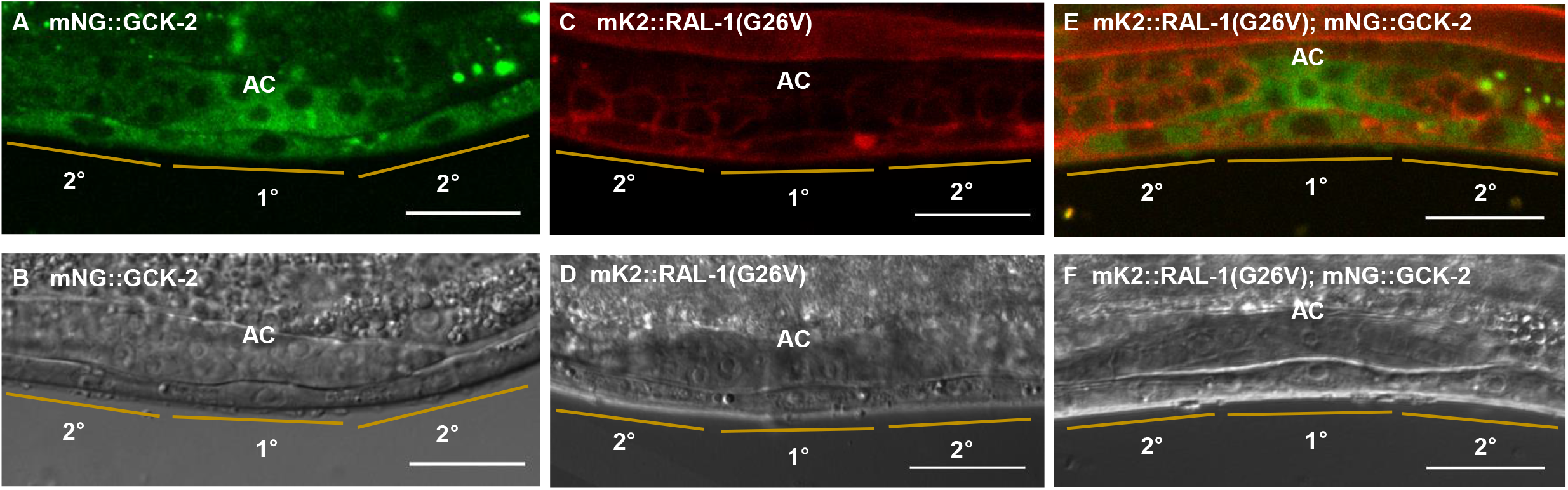
Endogenously tagged GCK-2 and RAL-1 are expressed in VPCs. Representative confocal and DIC micrographs of the presumptive 1° (P6.p) and 2° (P5,7.p) VPCs of *gck-2(re113[mNeonGreen^3xFlag::gck-2])* animals. (C, D) Confocal and DIC micrographs of the presumptive 1° (P6.p) and 2° (P5,7.p) VPCs of *ral-1(re160gf[mKate2^3xFlag::ral-1(G26V)])* animals. (E, F) Merged confocal and DIC micrographs of the presumptive 1° and 2° VPCs of the *ral-1(re160gf); gck-2(re113)* double mutant, from a separate animal than the single tags. “AC” label is placed directly above the Anchor Cell. Scale bar = 20 μm.

Critically, in light of complex results from *exoc-8(ok2523)* in different backgrounds (Fig. S2D; see above), *rels10[ral-1(gf)]-dependent* ectopic 2° induction was blocked by *exoc-8(ok2523)* (Fig. 5B). That RAL-1 2°-promoting activity depends on EXOC-8 is consistent with EXOC-8 functioning downstream of RAL-1 to transduce the 2°-promoting signal. We therefore speculate that a RAL-1-EXOC-8-GCK-2 cascade transduces a 2°-promoting signal, while acknowledging that EXOC-8 may perform other functions (see Discussion).

Multiple lines of evidence indicate that LIN-15 and many other genes in the “synMuv” group cooperate to redundantly restrict LIN-3/EGF expression to the AC (Cui et al., 2006; Fay and Yochem, 2007; Herman and Hedgecock, 1990; Huang et al., 1994; Myers and Greenwald, 2005). We previously exploited this feature of the vulval system to titrate LIN-3/EGF “dose” to induce ectopic 2° but not 1° VPCs in the *lin-12(n379*d) background (Zand et al., 2011), consistent with earlier manipulations of LIN-3/EGF and LET-23/EGFR signals to promote 2° fate, which supported the Morphogen Gradient Model (Katz et al., 1995; Katz et al., 1996). At 15°C, a temperature-sensitive mutation in *lin-15, n765ts*, supports normal vulva induction without inducing ectopic 1 ° cells. But in the *lin-12(n379*d) background at 15°C, n765ts strongly increased ectopic 2° induction. We showed that this 2°-promoting activity depends on LIN-3/EGF, LET-60/Ras, RGL-1/RalGEF, and RAL-1: RNAi depletion of *let-60, rgl-1*, and *ral-1* blocked the increased ectopic 2°s conferred by *lin-12(n379d); lin-15(n765ts)* at 15°C. We further showed that excess expression of LIN-3/EGF and an activating mutation in LET-23/EGFR conferred similar promotion of 2° fate via activation of the LET-60-RGL-1-RAL-1 module (Zand et al., 2011).

As expected, *ral-1(gk628801*rf) decreased ectopic 2° induction in the *lin-12(n379d); lin-15(n765ts)* background at 15°C (Fig. 5C), validating our prior results using *ral-1(RNAi)* (Zand et al., 2011). Similarly, *gck-2(RNAi)* and *gck-2(ok2867)* reduced ectopic 2° induction in the *lin-12(n379d); lin-15(n765ts)* background (Figs. 5D; S5A). Thus, we conclude that the 2°-promoting signal of EGF is, at least in part, GCK-2-dependent, consistent with RAL-1 signaling though GCK-2.

### Kinase-dependent GCK-2 functions cell autonomously in VPCs

Our fourth criterion for a RAL-1 effector is that its 2°-promoting activity function cell autonomously. We generated transgenic extrachromosomal arrays expressing VPC-specific GCK-2(+) and assessed rescue of mutant *gck-2* suppression of activated RAL-1. In the *rels10[ral-1(gf)]; lin-12(n379d); gck-2(tm2537)* and *rels10[ral-1(gf)]; lin-12(n379d); gck-2(ok2867)* backgrounds, VPC-specific expression of wild-type GCK-2 restored the increased 2° induction phenotype suppressed by *gck-2* mutations (Figs. 5E; S5B; S5C). Conversely, in the same backgrounds VPC-specific expression of putative kinase dead (KD) GCK-2 (for HPK1/MAP4K1, “GCK-2 group”; Kiefer et al., 1996) failed to rescue mutant *gck-2* suppression of *rels10* (Figs. 5F; S5D). Remember that *gck-2(ok2867)* enhanced ectopic 1° induction in the *let-60(n1046*gf) background (see Fig. 3E, above). VPC-specific expression of GCK-2(+) had no effect in the *let-60(n1046*gf) background, controlling for effects of VPC-specific GCK-2(+) overexpression (Fig. S5E). VPC-specific expression of GCK-2(+) restored baseline levels of ectopic 1° induction in the *let-60(n1046gf); gck-2(ok2867)* background (Fig. S5E). Taken together, these results suggest that GCK-2 functions cell autonomously in VPCs, via its kinase activity.

### RAL-1 and GCK-2 are expressed in VPCs

Our fifth criterion is that a RAL-1 effector be expressed in the VPCs. When analyzing the *sEx10525*[P*_gck-2_::gfp*+*dpy-5*(+)] transcriptional reporter transgene (Hunt-Newbury et al., 2007), we observed no GFP expression in VPCs. Therefore, we used CRISPR-Cas9 genome editing to insert mNG::3xFlag into the 5’ end of the endogenous *gck-2* gene, generating *gck-2(re113[mNG^3xFlag::gck-2])* (Fig. S6A). We confirmed alleles by western blot (Fig. S6B). We observed cytosolic tagged GCK-2 throughout vulval development (Figs. 6A, 6B; S6C-F). We observed the expression of GCK-2 in all tissues, including the germline and embryos in the adult hermaphrodite (Figs. S6G, S6H).

We also inserted mKate2::3xFlag into the 5’ end of the endogenous *ral-1* gene, with and without the activating G26V mutation (Figs. S6I, S6J, S6Q, S6R). The observed subcellular localization of RAL-1 and RAL-1(G26V) was similar, suggesting that the activating mutation does not alter localization. We observed tagged RAL-1 and RAL-1(G26V) throughout vulval development (Figs. 6C, 6D; S6K-N; S6S-X). Tagged RAL-1 and RAL-1(G26V) expression in the animal was ubiquitous, including vulva and germline, and localized primarily to plasma membrane and adherens junctions (Figs. S6O, S6P, S6Y, S6Z). To assess co-localization of RAL-1 and GCK-2, we made the *ral-1(re160gf[mKate2^3xFlag::ral-1(G26V)]); gck-2(re113[mNG^3xFlag::gck-2])* strain. We observed strong localization of RAL-1(G26V) to plasma membrane and junctions and GCK-2 to cytosol at Pn.p (1-cell) and Pn.px (2-cell) stages (Figs. 6E, 6F; S6AA, S6AB). Thus, expression of tagged endogenous RAL-1 (wild-type and G26V) and GCK-2 are expressed in VPCs, one of our criteria for a RAL-1 -GCK-2 signaling cascade. However, we did not observe evidence of 2°-specific recruitment of GCK-2 to the plasma membrane by activated RAL-1. We will consider this incongruity further in the Discussion.

### The PMK-1/p38 MAP kinase functions downstream of RAL-1 and is expressed in VPCs

The “GCK-2 group” is part of the Ste20 family of MAP4 kinases and is frequently associated with activation of JNK or p38 MAP kinase cascades (Dan et al., 2001; Delpire, 2009). Based on our model that RAL-1 signals through GCK-2, we investigated components of MAPK cascades as functioning downstream of RAL-1, identifying MLK-1/MAP3K and PMK-1/p38 as putative components of the RAL-1-GCK-2 2°-promoting signaling cascade.

*C. elegans* encodes orthologs of MAP3Ks and MAP2Ks (Sakaguchi et al., 2004). The *km19* deletion in MLK-1/MLK/MAP3K (Mizuno et al., 2004) enhanced *let-60(n1046*gf) vs. *n1046* alone (p=0.009, N of 90 and 60, respectively), consistent with MLK-1/MAP3K acting in this cascade. In contrast, the *ok1382* deletion in MTK-1/MEKK4/MAP3K failed to enhance *let-60(n1046*gf) vs. *n1046* alone (p=0.4, N of 50 and 91, respectively). The *km4* deletion in SEK-1/MKK3/6/MAP2K (Tanaka-Hino et al., 2002) and the *ok1545* deletion in MKK-4/MKK4/MAP2K failed to enhance *let-60(n1046*gf) vs. *n1046* alone (p=0.6 N of 60, 60; and p=0.2 N of 90, 90, respectively). Several other MAP2Ks were not tested.

*C. elegans* encodes three p38/MAPK paralogs in an operon, in order: PMK-2, PMK-3, and PMK-1. Of these, PMK-2 and PMK3 are expressed primarily in intestine, while PMK-1 is expressed more broadly. All three are thought to contribute to innate immunity and stress response (Mertenskotter et al., 2013). Fitting some of our criteria for a RAL-1 effector, putative null *pmk-1(km25)* (Mizuno et al., 2004) conferred increased ectopic 1° induction with *let-60(n1046*gf) and blocked increased ectopic 2° induction with ral-1(gf); *lin-12(n379*d) (Figs. 7A, 7B).

We tagged the endogenous *pmk-1* gene with mNG::3xFlag at the 3’ end (Figs. S7A, S7B). We observed PMK-1::mNG expression in vulval lineages and throughout the rest of the animal, with the exception of the germline; germline expression appeared to be silenced, an established phenomenon with certain foreign DNA insertions (Fig. 7C, 7D; S7C-F; Dickinson et al., 2015). Thus, expression of PMK-1 meets the criteria of an effector that functions downstream of RAL-1. Using a transgenic pmk-1 promoter translational GFP fusion *(Ppmk-1::pmk-1::gfp;* Mertenskotter et al., 2013), we observed no vulval signal (Figs. S7G-J). However, since *pmk-1* is the last gene in an operon, key regulatory elements may be absent from the construct.

**Figure 7.**
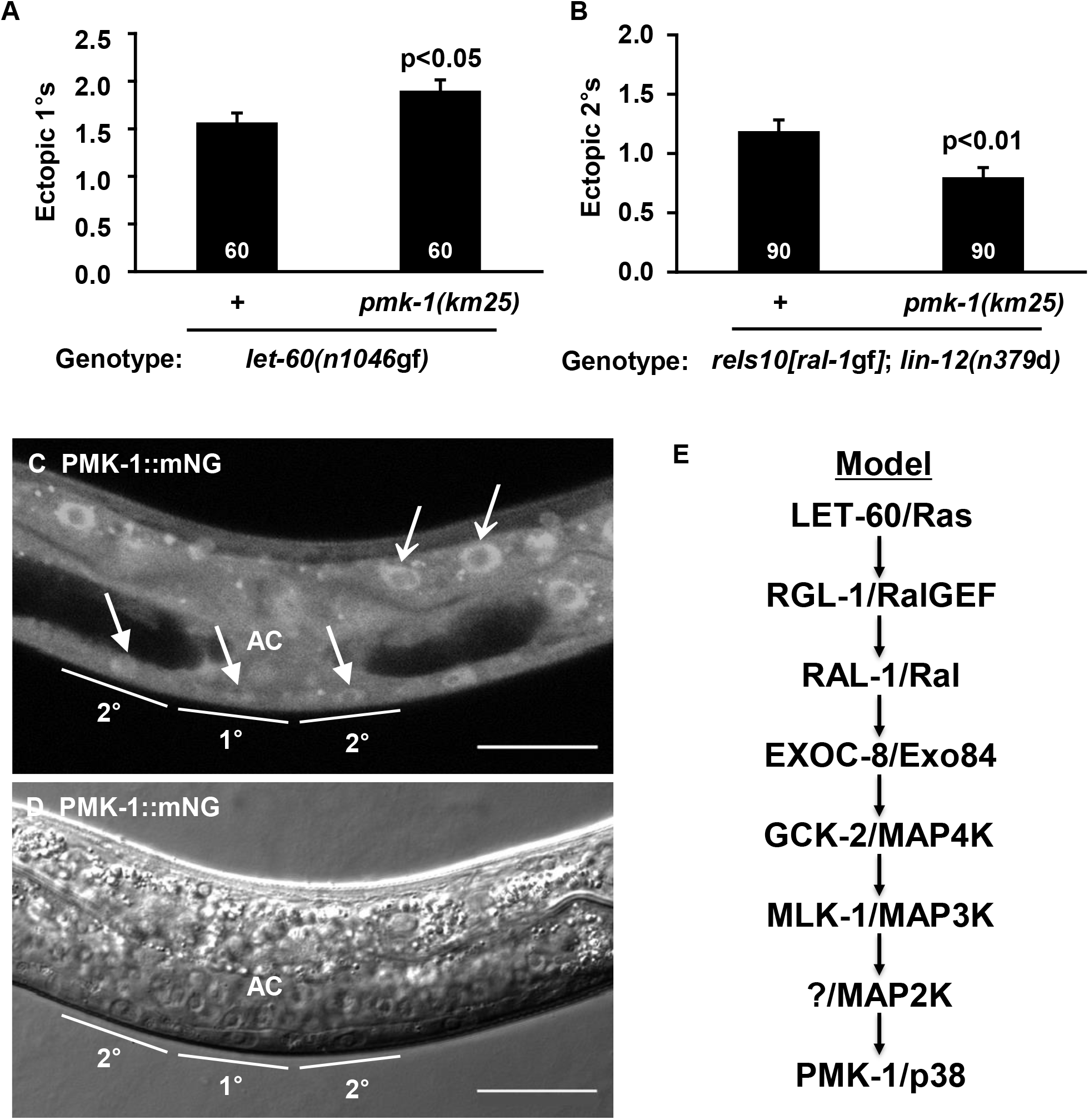
PMK-1/p38 functions downstream of RAL-1. (A) Putative null, *pmk-1(km25)* confers increased ectopic 1°s in the *let-60(n1046*gf) background. (B) *pmk-1(km25)* blocks the increased 2°s in the *rels10[P_lin_-31::ral-1(gf)]; lin-12(n379*d) background. P values calculated by *t* test. Error bars = S.E.M. (C, D) Representative confocal and DIC micrographs of *pmk-1(re170[pmk-1::mNG^3xFlag])*. Endogenously tagged PMK-1 is expressed in VPCs. Solid arrow: VPC nuclei. Open arrow: Gut nuclei. Scale bar = 20 μm. (F) A signaling transduction model. A LET-60/Ras-RGL-1/RalGEF-RAL-1/Ral-EXOC-8/Exo84-GCK-2/MAP4K-MLK-1/MAP3K-PMK-1/MAPK cascade promotes 2° fate in *C. elegans* VPC fate patterning.

We had hoped to observe activity-dependent cytosol-to-nucleus translocation (Ben-Levy et al., 1998) of PMK-1 as a biomarker for upstream RAL-1 signaling activation. Instead, we observed nuclear and cytosolic localization in all cells, including vulval lineages (Figs. 7C, 7D; S7C-F). We considered our C-terminal CRISPR tagging scheme may have altered PMK-1 localization, even though we did not observe phenotypic changes conferred in the sensitized background by PMK-1::mNG (see Star Methods). Therefore, we N-terminally tagged PMK-1 expressed from *lin-31* promoter in VPCs, using the miniMos system (de la Cova et al., 2017), and with mKate2 rather than mNG. We observed localization to both nuclei and cytosol of VPCs (Figs. S7K-N). Consequently, we propose that our tagging strategies do not disrupt PMK-1 function, but rather that part of the endogenous PMK-1 population is constitutively targeted to the nucleus. We speculate that over-expression from the extrachromosomal array shows mostly cytosolic localization because only a small subset of the total PMK-1 molecules occupy the nucleus, and that the proportion of nuclear PMK-1 to total protein is much higher when looking at endogenous rather than over-expressed protein.

## Discussion

We found that EXOC-8/Exo84 contributes to the LET-60/Ras-RGL-1/RalGEF-RAL-1/Ral 2°-promoting signal during patterning of *C. elegans* VPC fate. By our genetic criteria GCK-2, a CNH domain-containing MAP4K orthologous to *Drosophila* Hppy and mammalian MAP4K1, 2, 3 and 5, is a downstream effector of RAL-1, as are MLK-1/MAP3K and PMK-1/p38 MAPK (Fig. 7E). The LET-60-RGL-1 -RAL-1 -EXOC-8-GCK-2-MLK-1 -PMK-1 cascade is a nonessential 2°-promoting signal in *C. elegans* VPC fate patterning that supports the essential 2°-promoting signal via LIN-12/Notch. We showed that GCK-2 and PMK-1 function downstream of RAL-1 cell autonomously, and that GCK-2 is sufficient to promote 2° fate in support of LIN-12/Notch. We did not observe evidence of ectopic 2° induction by *rels10[ral-1(gf)]* or *ral-1(re160*gf). Thus, given the modest modulatory role of the RAL-1 2°-promoting cascade, there is no reason to propose that RAL-1 is sufficient to induce 2° fate, though the actual experiment in the absence of *lin-12* is as yet prohibitively difficult (abrogation of LIN-12 function duplicates the anchor cell, and thus results in complex VPC induction; Greenwald et al., 1983).

Sec5 and Exo84 are subunits of the exocyst complex and known Ral binding partners in mammals (reviewed in Gentry et al., 2014; Kashatus, 2013). Ral-Exo84 and Ral-Sec5 regulate exocytosis, cancer cell proliferation, and immunity (Chien et al., 2006; Fukai et al., 2003; Issaq et al., 2010; Jin et al., 2005; Moskalenko et al., 2002; Moskalenko et al., 2003; Sugihara et al., 2002). Yet we do not understand how Ral signaling is propagated through the Sec5 and Exo84 exocyst partners. Exo84 and Sec5 also confound biochemical identification of downstream signaling partners, since the exocyst is involved in central cell biological processes and potentially interacts with myriad partners (Tanaka et al., 2017; Wu and Guo, 2015). There are some exceptions: RalB-Sec5 directly recruits and activates the atypical IκB kinase family member TBK1 to contribute to human cancer cell survival (Chien et al., 2006), RalB-Exo84 promotes autophagosome assembly under starvation conditions in human epithelial cells (Bodemann et al., 2011), while under replete conditions RalB-Sec5 stimulates mTORC1 activation in pancreatic tumor cells to promote cell invasion and inhibit autophagy (Martin et al., 2014). Yet we lack biomarkers for activated Ral and have limited knowledge of effectors downstream of Sec5 and Exo84. Using developmental patterning of the *C. elegans* VPCs as a simple model system, we defined a potentially new signaling cascade downstream of RAL-1 in development. Additional signaling cascades may function downstream of RAL-1 in different tissues.

The GCK-2 paralog, MIG-15, also contributes to VPC patterning: depletion of *mig-15* derepressed both 2°- and 1°-inducing backgrounds, respectively *lin-12(n379*d) and *let-60(n1046*gf). A 3°-promoting gene might be predicted to similarly antagonize both 1°- and 2°-promoting signals. However, the connection of MIG-15 to 1°-promoting signals and 2°-promoting signals is unclear, and is complicated by the role of MIG-15 in vulval morphogenesis. In contrast, we clearly delineated GCK-2 genetically as a component of a positive regulatory cascade downstream of RAL-1-EXOC-8.

While mutation of neither RLBP-1/RalBP1 nor SEC-5/Sec5 altered vulval patterning in sensitized backgrounds, mutation of EXOC-8/Exo84 conferred phenotypes consistent with functioning as a signaling intermediary in a RAL-1 2°-promoting cascade. However, mutation of EXOC-8 also conferred defects consistent with other activities in VPC fate patterning: unlike reduced RAL-1 or GCK-2 function, reduced EXOC-8 function conferred increased 2° induction in the *lin-12(n379*d) 2°-inducing background. We speculate that EXOC-8 performs at least two functions: (1) an intermediary in RAL-1-GCK-2 2°-promoting signaling, and (2) in an anti-2° capacity, perhaps with MIG-15. Thus, the roles of EXOC-8 and MIG-15 in VPC fate patterning are enigmatic, and will be the subject of future genetic and biochemical studies, particularly as we develop better experimental tools via use of CRISPR.

Activation of the p38 and JNK families of MAPKs have variously been associated with activation of the GCK-I (GCK-2) and GCK-IV (MIG-15) subfamilies of CNH domain MAP4Ks (Delpire, 2009). Neither subfamily has been studied systematically. Our ongoing observation of the literature is consistent with the GCK-2 and MIG-15 subfamilies generally being associated with p38 and JNK, activation, respectively. Yet so many of these studies depend on protein over-expression that we hesitate to draw general conclusions. Here we connect GCK-2 with PMK-1/p38 function, while in *Drosophila* dorsal closure of the embryo Msn (MIG-15 subfamily) is associated with JNK function (Su et al., 2000; Su et al., 1998).

Activation of MAP kinases is often associated with cytosol-to-nuclear translocation, initially shown with the canonical ERK MAPK (Gonzalez et al., 1993; Lenormand et al., 1993) but also shown for p38 MAPK (Ben-Levy et al., 1998). Extrachromosomal transgenic *C. elegans* PMK-1::GFP similarly translocated from cytosol to the nucleus upon stress (Mertenskotter et al., 2013). Yet we found that this PMK-1::GFP fusion was not expressed in the VPCs, perhaps because *pmk-1* is expressed as part of a multi-gene operon, and so the transgene is missing key regulatory sequences. Further, the transgenic PMK-1::GFP is likely to be over-expressed, as is typical for *C. elegans* transgenic arrays, and thus may obscure more nuanced regulatory inputs. Consequently, we generated endogenous PMK-1::mNG via CRISPR and also introduced VPC-specific single-to-low copy mK2::PMK-1 (Fig. 7, Fig S7). While we observed that endogenous PMK-1::mNG was expressed in VPCs, to our surprise we observed consistent nuclear PMK-1::mNG throughout the animal, including VPCs throughout their fate patterning, and nuclear mK2::PMK-1 in VPCs throughout their patterning. One could speculate that we perturbed PMK-1 function both through C- and N-terminal tagging, yet we showed that PMK-1::mNG function appeared normal in the *let-23(sa62*gf) background. These observations leave us at an impasse. Is the prevailing model for MAP kinase activation flawed? Is PMK-1/p38, a known stress kinase, tonically activated as a consequence of endogenous stressors or culture conditions? Or is translocation a modest part of the activation process that was previously masked by assay conditions? Our observation may lead to important mechanistic considerations of p38 activation and activation of MAP kinases in general, and is worth further investigation.

Many small GTPases, including mammalian RalA and RalB, are membrane-targeted through prenylation. Based on its C-terminal CAAX sequence, RAL-1 is inferred to be geranylgeranylated (Reiner and Lundquist, 2016), and our CRISPR tag of endogenous RAL-1 showed strong localization to the plasma membrane in all cells. The canonical mechanism for effector activation, originally defined for Ras-Raf (Block et al., 1996; Chiu et al., 2002), is recruitment of cytosolic effector to the plasma membrane by activated small GTPase. While our genetic analysis indicates that GCK-2 functions downstream of the 2°-promoting RAL-1 in VPCs, we did not observe co-localization of tagged endogenous RAL-1 and GCK-2, or activity-dependent enrichment of plasma membrane GCK-2, in presumptive 2° cells. This observation could be explained by activated, membrane-tethered RAL-1 recruiting only a small portion of cytosolic GCK-2. Alternatively, perhaps RAL-1 effector activation proceeds through an atypical mechanism, consistent with the non-canonical nature of the exocyst as an effector. The exocyst presents an interesting conundrum in signaling: it is clearly required for much Ral signaling (reviewed in Gentry et al., 2014), yet thwarts conventional biochemical bootstrapping through signaling cascades. Thus, the lack of co-localization of RAL-1 and GCK-2 could also be explained by certain populations of RAL-1 and GCK-2 being constitutively associated, perhaps at the exocyst complex. Though not the same subfamily (GCK-2 vs. MIG-15), this model is consistent with co-immunoprecipitation of mammalian Sec5 and HGK/MAP4K4 (Balakireva et al., 2006). We speculate that the complex of RAL-1/EXOC-8/GCK-2 recruits a co-activator when RAL-1 is activated. Such a mechanism was implicated in studies of mammalian MAP4K3 (in the GCK-2 subfamily), in which a putative activating phosphorylation event was detected (Yan et al., 2010). While shown to be inhibited by PP2AT61 ε, the kinase(s) mediating this phosphorylation event remains unknown. Alternatively, perhaps RAL-1 and GCK-2 never physically interact, and thus genetic analysis was required to reveal this cascade.

The advent of CRISPR-based tools permits analysis of endogenous proteins, and hence potentially improved cell biological and biochemical analysis. For example, all known Ral binding partners were discovered by yeast two-hybrid analysis, an approach with a strong record but also ample false negatives, say, in conditions of activity-dependent interactions or metazoan-specific subcellular localization. Thus, we may be able to use biochemical approaches to with tagged endogenous RAL-1 to identify novel interactors. Yet we also recognize the balance of strengths and weaknesses in the model invertebrate system, so mammalian cell-based studies may complement our genetic analysis to elucidate details of molecular mechanisms.

It is as yet unclear whether our findings of a genetic requirement for PMK-1 downstream of the RAL-1 2°-promoting activity herald a similar use of p38 downstream of mammalian Ral isoforms during development or cancer. An alternative possibility is that Ral in various metazoa signals through an array of effectors, only of some of which are relevant to cancer. Genetic and tissue heterogeneity of tumors, coupled with the historic difficulty of assessing Ral activation levels in tumors, make this question non-trivial to address rigorously. This question is further complicated by the diverging functions of RalA and RalB in cancer, and their mostly poorly defined roles in non-pathogenic development and physiology. Yet this study could lead to surveys of phospho-p38 levels in Ras-positive tumors as a function of RalA or RalB, thus potentially satisfying the great demand for cancer biomarkers of Ral activation. Alternatively, establishment of a viable *in vivo* invertebrate model for RAL-1 function, added to the development of elegant genetic tools, should lead to extensive investigation of RAL-1 in diverse areas of biology, potentially leading us to clinically important biomarkers and “druggable targets” other than p38.

In conclusion, we have demonstrated that the LET-60/Ras-RGL-1/RalGEF-RAL-1/Ral 2°-promoting signal acts through exocyst component EXOC-8/Exo84 to trigger a GCK-2/MAP4K-PMK-1/p38 cascade. From a developmental biology standpoint, the mechanism by which the EGF morphogen gradient promotes 2° VPC fate in support of LIN-12/Notch is of long-standing interest (Kenyon, 1995). From a cancer biology standpoint, this study may also contribute to development of diagnostic biomarkers and small molecule inhibitors for Ras- and Ral-dependent cancers.

## STAR METHODS

Detailed methods are provided in the online version of this paper and include the following:

- KEY RESOURCES TABLE
- CONTACT FOR REAGENT AND RESOURCE SHARING
- EXPERIMENTAL MODEL AND SUBJECT DETAILS

- *C. elegans* Strains
- METHOD DETAILS

- Plasmids and Transgenes Used for Expression in *C. elegans*
- CRISPR design
- Image Acquisition
- QUANTIFICATION AND STATISTICAL ANALYSIS
- DATA AND SOFTWARE AVAILABILITY

**STAR METHODS Table 1.** Alleles

Alleles
| Allele | Lesion | Functional interpretation |
| --- | --- | --- |
| *let-60(n1046gf)* | G13E | Moderate gain of function |
| *lin-12(n379d)* | E889K | Weak gain of function |
| *lin-3(n378rf)* | E21K | Reduced function |
| *ral-1(gk628801)* | R139H | Reduced function |
| *exoc-8(ok2523)* | 1474 bp out-of-frame deletion | Putative null |
| *sec-5(pk2357)* | M389* nonsense | Strong loss of putative null |
| *exoc-7(ok2006)* | 1803 bp out-of-frame deletion | Putative null |
| *rlbp-1(tm3665)* | 346 bp out-of-frame deletion | Putative null |
| *gck-2(ok2867)* | 549 bp in-frame deletion | Kinase dead |
| *gck-2(tm2537)* | 565 bp out-of-frame deletion | Putative null |
| *mig-15(rh148)* | V168E | Moderate reduction |
| *mig-15(rh80)* | W898* nonsense | Strong reduction, not null |
| *lin-15A/B(n765ts)* | 171 bp deletion/982 bp insertion | ts loss of function |
| *pmk-1(km25)* | 375 bp out-of-frame deletion | Putative null |

**STAR METHODS Table 2.**
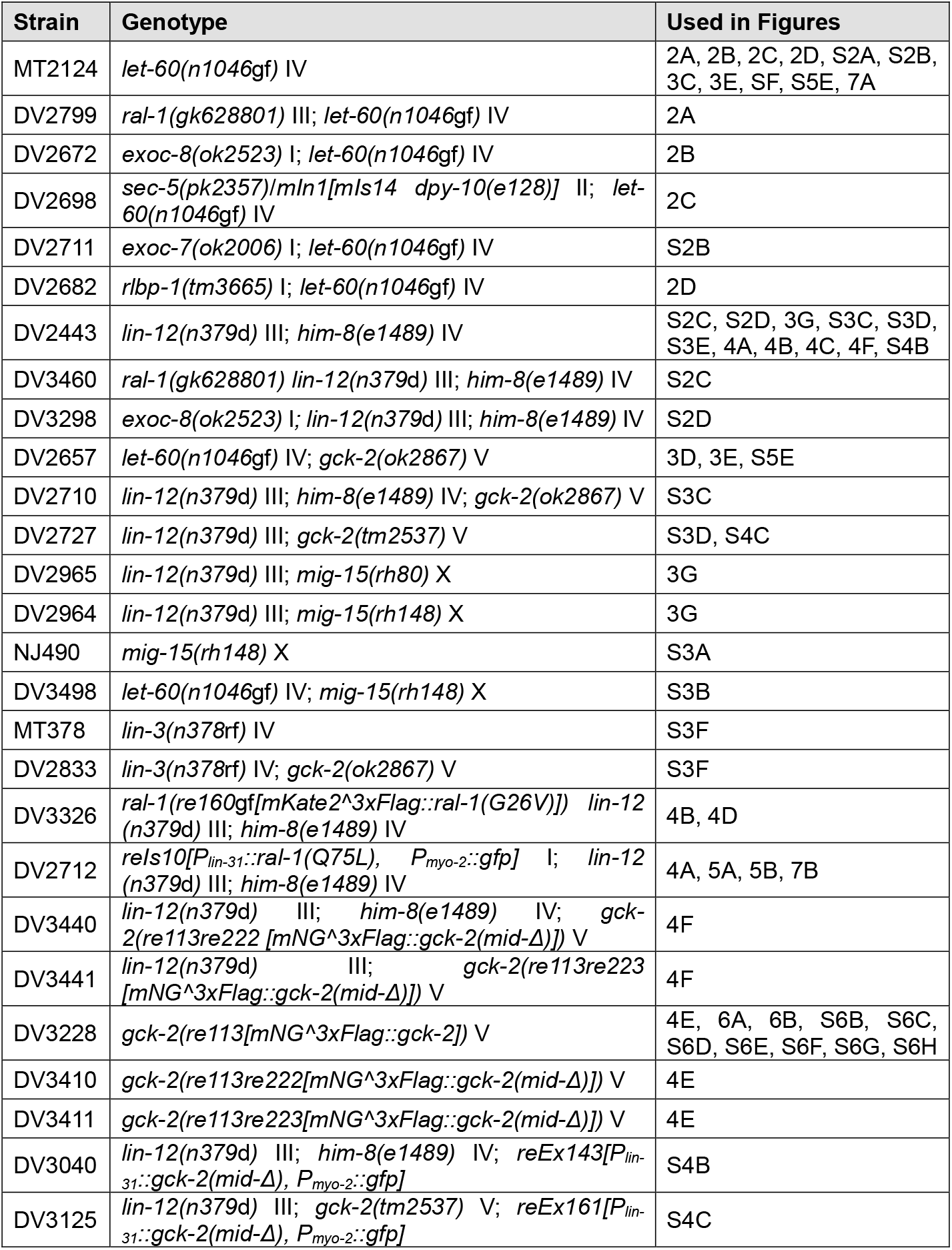
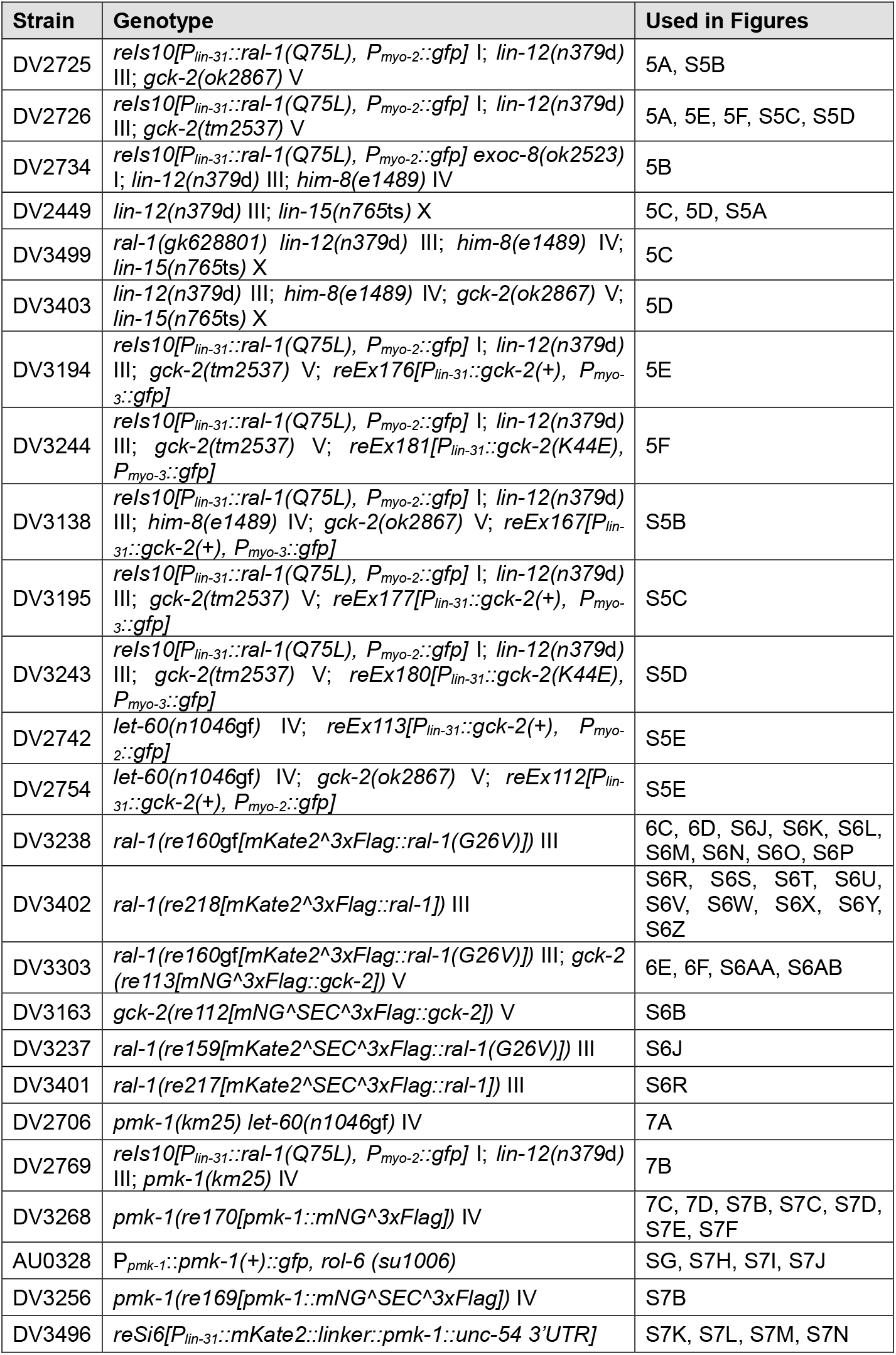
Strains

**STAR METHODS Table 3.**
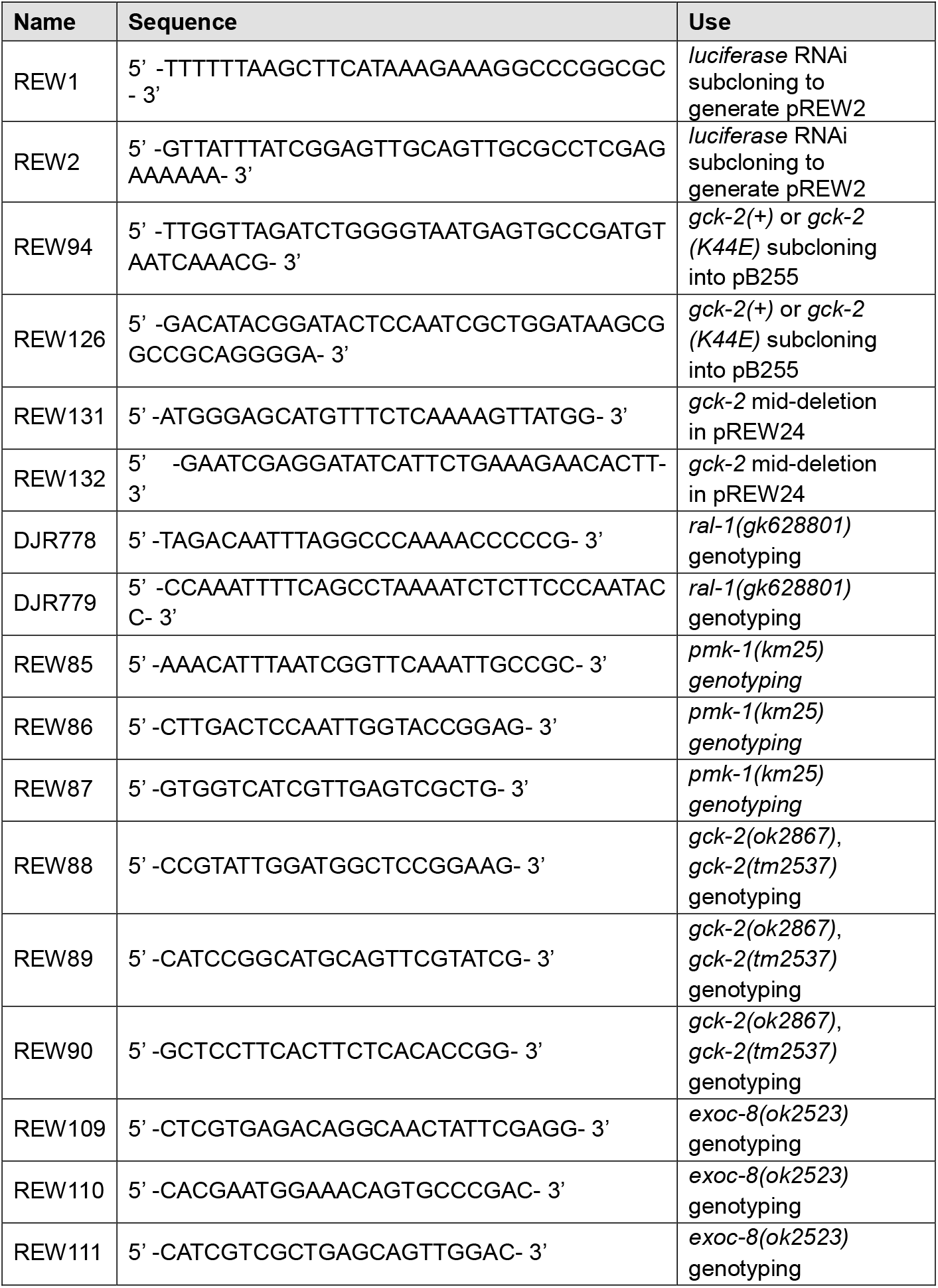
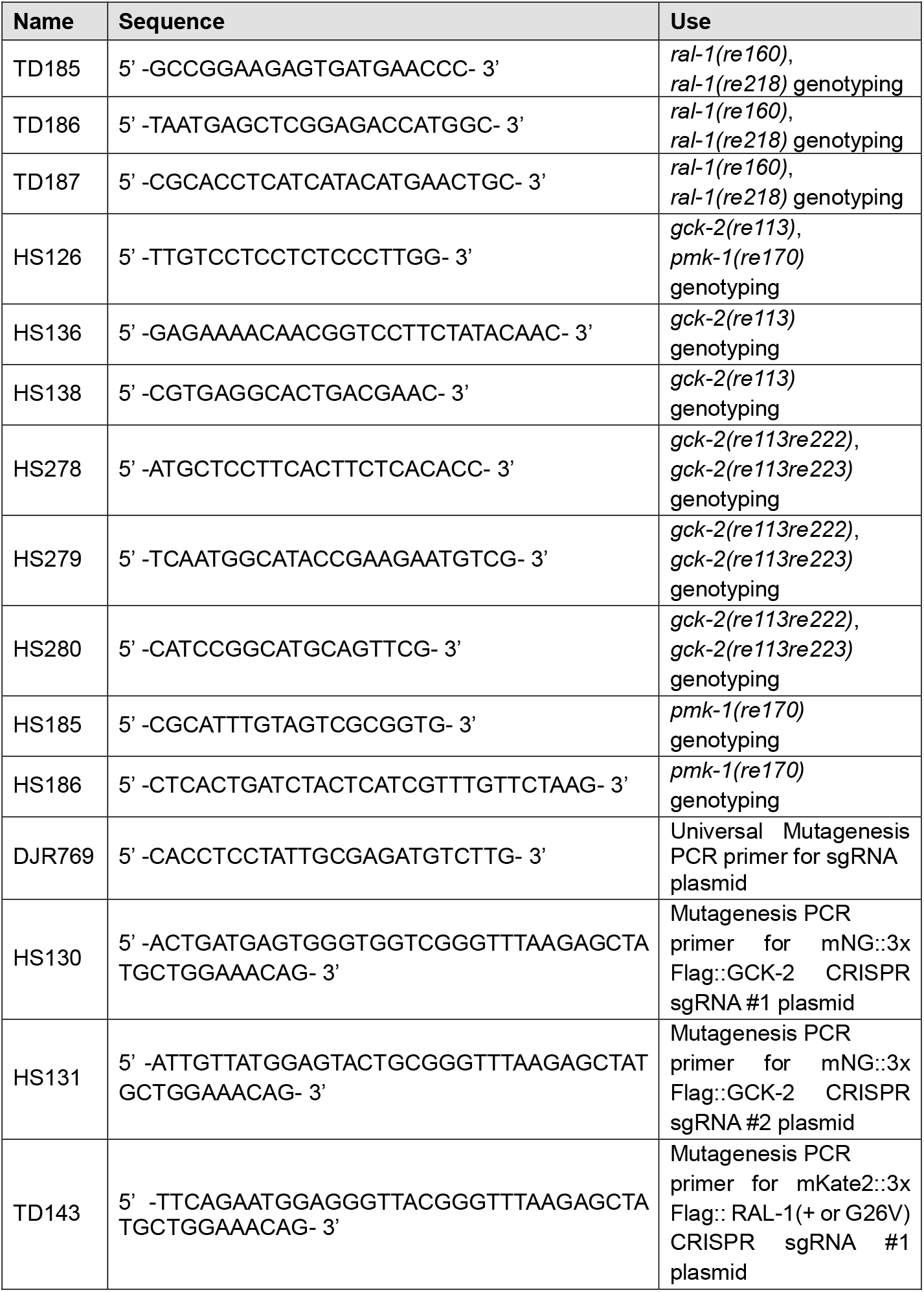
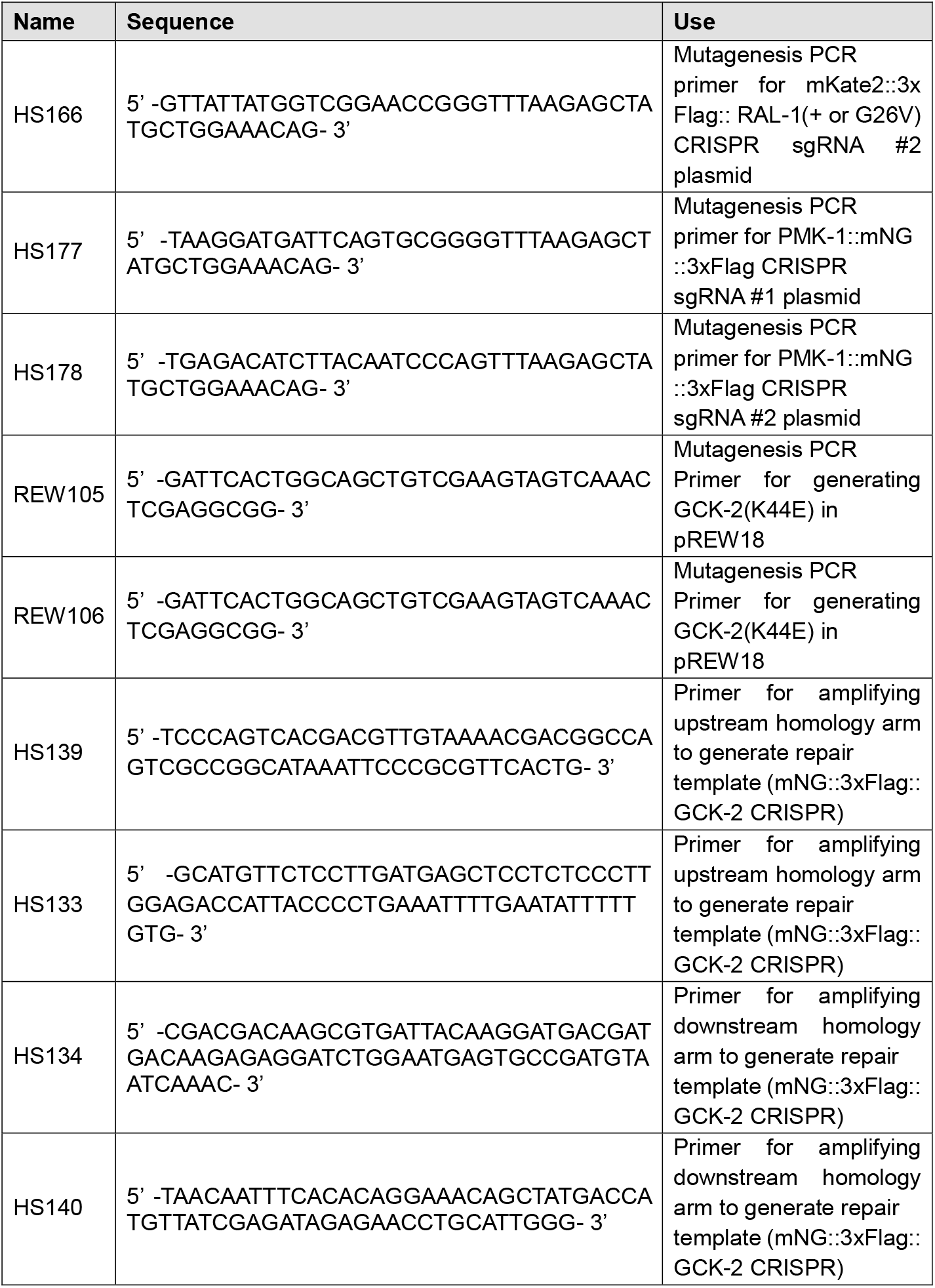
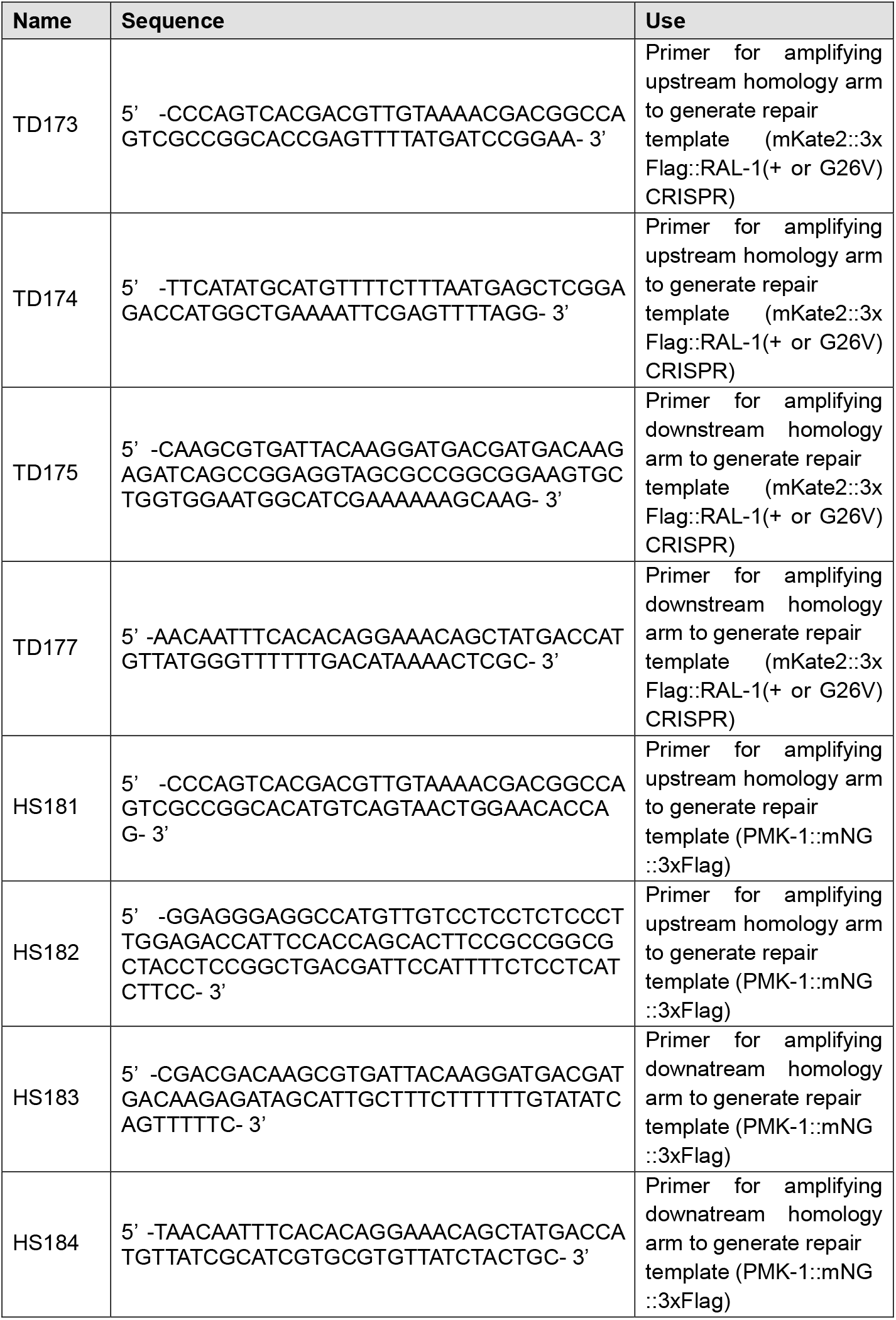
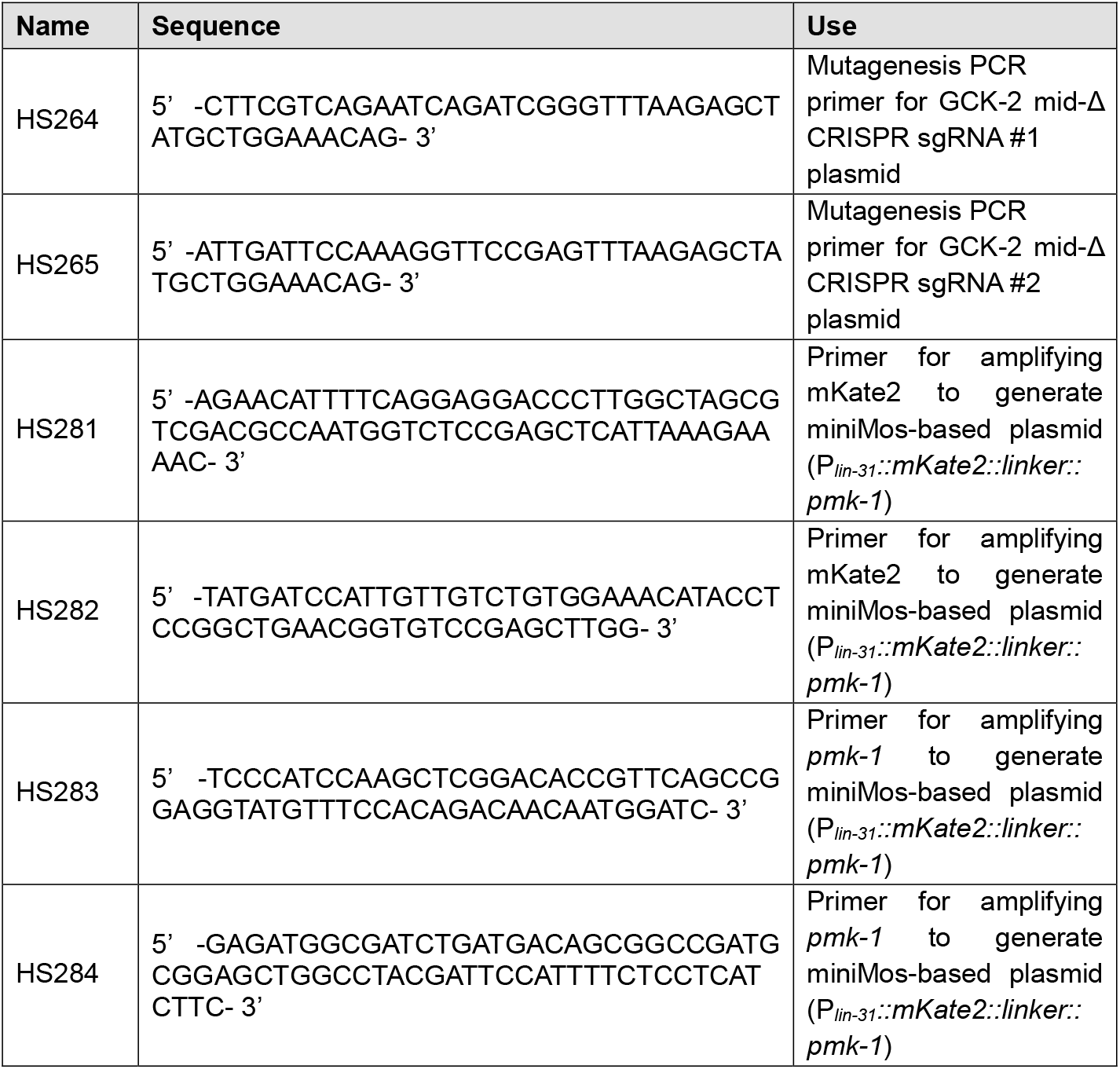
Primers

**STAR METHODS Table 4.** Plasmids

| Name | Description | Used for |
| --- | --- | --- |
| pREW2 | RNAi of *luciferase* was subcloned into L4440 | Negative control for RNAi experiments |
| pREW18 | *P<sub>lin-31</sub>::gck-2(+):let-858* 3'UTR | Rescue experiment in *let-60(n1046gf)*; *gck-2(ok2867)* |
| pREW24 | *P<sub>lin-31</sub>::gck-2(+):unc-54* 3'UTR | Rescue experiment in *rels10[ral-1(gf)]*; *lin-12(n379d)*; *gck-2(ok2867)* |
| pREW25 | *P<sub>lin-31</sub>::gck-2(mid-deletion):unc-54* 3'UTR | 2° induction experiment in *lin-12(n379d)* and *lin-12(n379d)*; *gck-2(tm2537)* |
| pREW32 | *P<sub>lin-31</sub>::gck-2(K44E):unc-54* 3'UTR | Kinase dead negative control rescue experiment in *rels10[ral-1(gf)]*; *lin-12(n379d)*; *gck-2(ok2867)* |
| pHS21 | Homology arms (534 bp upstream and 875 bp downstream) was subcloned into pDD268 | mNG<sup>3xFlag</sup>::GCK-2 CRISPR knock-in (repair template) |
| pHS13 | Mutagenized pJW1236 using primers, DJR769 and HS130 | mNG<sup>3xFlag</sup>::GCK-2 CRISPR knock-in (sgRNA-Cas9 plasmid) (sgRNA #1) |
| pHS14 | Mutagenized pJW1236 using primers, DJR769 and HS131 | mNG<sup>3xFlag</sup>::GCK-2 CRISPR knock-in (sgRNA-Cas9 plasmid) (sgRNA #2) |
| pTD36 | Homology arms (869 bp upstream and 754 bp downstream) was subcloned into pDD285 | mKate2<sup>3xFlag</sup>::RAL-1(+) CRISPR knock-in (repair template) |
| pTD40 | Homology arms (869 bp upstream and 754 bp downstream) was subcloned into pDD285 | mKate2<sup>3xFlag</sup>::RAL-1(G26V) CRISPR knock-in (repair template) |
| pTD38 | Mutagenized pJW1236 using primers, DJR769 and TD143 | mKate2<sup>3xFlag</sup>::RAL-1(+) and mKate2<sup>3xFlag</sup>::RAL-1(G26V) CRISPR knock-in (sgRNA-Cas9 plasmid) (sgRNA #1) |
| pHS23 | Mutagenized pJW1236 using primers, DJR769 and HS166 | mKate2<sup>3xFlag</sup>::RAL-1(+) and mKate2<sup>3xFlag</sup>::RAL-1(G26V) CRISPR knock-in (sgRNA-Cas9 plasmid) (sgRNA #2) |
| pHS40 | Homology arms (785 bp upstream and 875 bp downstream) was subcloned into pDD268 | PMK-1::mNG<sup>3xFlag</sup> CRISPR knock-in (repair template) |
| pHS26 | Mutagenized pJW1236 using primers, DJR769 and HS177 | PMK-1::mNG<sup>3xFlag</sup> CRISPR knock-in (sgRNA-Cas9 plasmid) (sgRNA #1) |
| pHS25 | Mutagenized pJW1236 using primers, DJR769 and HS178 | PMK-1::mNG<sup>3xFlag</sup> CRISPR knock-in (sgRNA-Cas9 plasmid) (sgRNA #2) |
| pHS38 | Mutagenized pJW1236 using primers, DJR769 and HS264 | GCK-2 mid-deletion CRISPR knock-in (sgRNA-Cas9 plasmid) (sgRNA #1) |
| pHS39 | Mutagenized pJW1236 using primers, DJR769 and HS265 | GCK-2 mid-deletion CRISPR knock-in (sgRNA-Cas9 plasmid) (sgRNA #2) |
| pHS41 | Sequence of *mKate2::linker::pmk-1* was subcloned into pCC249 (*P<sub>lin-31</sub>::unc-54* 3'UTR) | PMK-1 miniMos for generating *reSi6* [*P<sub>lin-31</sub>::mKate2::linker::pmk-1::unc-54* 3'UTR] |

**STAR METHODS Table 5.**
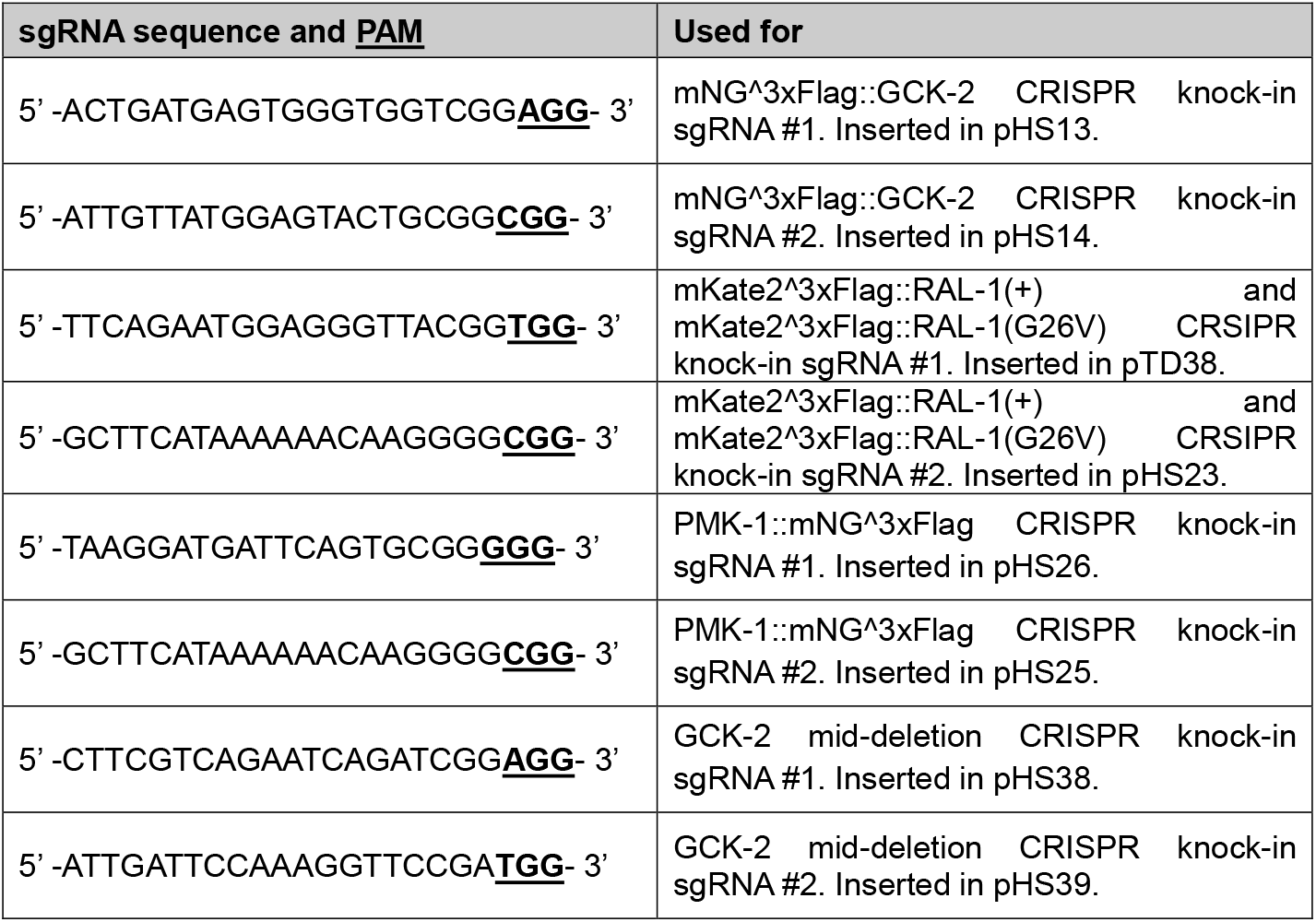
sgRNA sequences and <u>PAMs</u>

**STAR METHODS Table 6.**
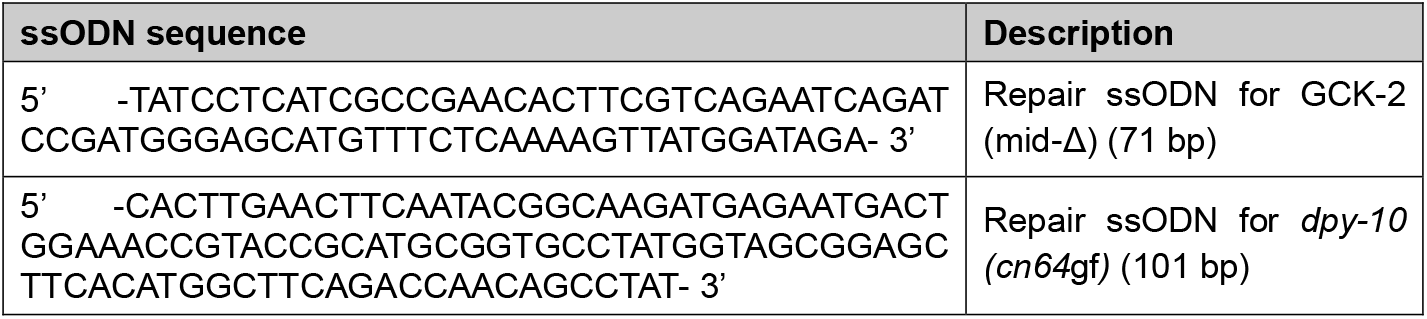
Repair ssODN

## Acknowledgments

We thank the knockout consortia (Mitani, Barstead and Moerman labs) plus I. Greenwald and N Kirienko for strains. We thank members of the Reiner lab for helpful discussions and critical reading of the manuscript. Some strains were provided by the CGC, which is funded by NIH Office of Research Infrastructure Programs (P40 OD010440). Wormbase was used constantly. We thank D. Dickinson and B. Goldstein (UNC) for sharing CRISPR SEC vectors prior to publication. We thank Dr. N. Kirienko (Rice) for sharing the AU0038 *pmk-1* reporter, and Drs. Kirienko and D. Garsin (UT Health) for sharing expertise about inflammatory response and *Comamonas sp*. bacteria. This work was supported by NIH grants GM085309 and GM121625 to D.J.R.

## Author Contributions

D.J.R., H.S. and R.E.W.K. conceived the study, D.J.R., H.S. and R.E.W.K. designed the experiments, H.S. and T.D. designed CRISPR tagging strategies, T.D. generated *ral-1(re160gf[mKate2^3xFlag::ral-1(G26V)])*, R.F. performed *mig-15(RNAi)* experiments, H.S and R.E.W.K. conducted the experiments, and H.S. and D.J.R. wrote the manuscript.

## STAR Methods

### *C. elegans* handling and genetics

All strains were derived from the N2 wild type. Nomenclature was as described (Horvitz et al., 1979). Animals were cultured using standard conditions on OP50 bacteria on NGM agar places at 20°C (Brenner, 1974) except where noted. Strains used are shown in Supplementary Table 1.

PCR primers are listed in Supplementary Table 2. Single animal genotyping PCR reactions used Taq PCR Master Mix (QIAGEN). Deletions in *gck-2* were detected by triplex primers REW88/89/90 (Tm: 58°C, 32 cycles), resulting in 298 bp (wild-type), 478 bp *(ok2867)*, and 462 bp *(tm2537)* amplicons. PCR products were sequenced to confirm reported allele perturbations (Khan et al. 2012). The *exoc-8(ok2523)* deletion was detected by triplex primers REW109/110/111 (Tm: 58°C, 32 cycles), resulting in 411 bp (wild-type) and 270 bp *(ok2523)* amplicons. The *pmk-1(km25)* was detected by PCR using triplex primers, REW85/86/87 (Tm: 55.5°C, 32 cycles), resulting in 345 bp and 591 bp (wild-type) and 216 bp *(km25). ral-1(gk628801)* was detected by primers DJR778/779 (Tm: 57°C, 32 cycles) to generate a 250 bp amplicon, followed by overnight digestion with HpyCH4IV (NEB) to yield wild-type (121 bp, 51 bp, 48 bp and 30 bp) and *gk628801* (151 bp, 51 bp and 48 bp) bands.

The *sec-5(pk2358)/+* mutation was maintained as a stable heterozygote by GFP-tagged balancer *mln1mls14*, and homozygotes were obtained by scoring non-green progeny.

### Vulval induction scoring assay

To score vulval induction, late L4 animals were mounted on slides with a 3% agar pad in M9 buffer with 5 mM sodium azide. Invaginations of ectopic pseudovulvae were scored under DIC/Nomarski optics (Nikon eclipse Ni). Images were captured using NIS-Elements AR 4.20.00 software. The vulval induction index for ectopic 1° and 2° induction was scored as described elsewhere. Briefly, we counted vulval invaginations, comprising cell lineages of single VPCs undergoing morphogenesis, which were distinct from the composite 2°-1°-2° normal vulva lineages (the “Christmas tree,” which we argue more closely resembles the Stanley Cup). As expected, in the *let-60(n1046*gf) background the normal vulva was oriented on the AC in the center of the gonad. The morphology of ectopic 1 ° lineages generally conformed with the symmetrical “cap” characteristic of isolated 1 ° lineages (Katz et al., 1995). In the *lin-12(n379*d) background, the AC and normal vulva were mostly absent, as described (Greenwald et al., 1983). The morphology of ectopic 2° lineages generally conformed with the asymmetric “beret” characteristic of isolated 2° lineages (Green et al., 2008; Katz et al., 1995). When the AC and normal vulva were present in a *lin-12(n379*d) animal, data from that animal were flagged and excluded from the final count of ectopic 2° cells.

As previously described (Zand et al., 2011), the *let-60(n1046*gf) strain is liable to drift, resulting in increased induction of 1 ° cells. We established many frozen strains of *n1046* single mutants, and *n1046* outcrossed to N2, and established that the typical baseline is 1.2 to 1.5 ectopic 1° cells. We have also consistently observed that the *n1046* baseline is increased by ~0.2 when grown on bacterially mediated RNAi, including *gfp* or *luciferase* control strains (Zand et al., 2011; this study). Consequently, for all strains harboring an *n1046* mutation, we use a stringent protocol to minimize drift: strains are scored and a parafilmed plate established immediately after construction or thawing, strains are refreshed (if necessary) by chunking, and animals are never grown for several generations in culture. Assays in which *n1046* control strains deviate from the expected baseline are discarded. When using this rigorous protocol, we rarely observe significant deviations from expected baselines. Each figure panel with VPC counts is from animals grown together at the same time.

### Plasmids, generation of transgenic lines and integrated lines

Details of plasmid construction are available upon request. Transgenic lines were generated by microinjection of pB255-derived plasmids (50 ng/μl) with co-injection marker (20 ng/μl; either pPD118.33 [P*_myo-2_::gfp*] or pPD93.97 *[P_myo-3_::gfp])* into the relevant strain and maintained by selecting for fluorescent animals. *rels10[P_lin-31_::ral-1(Q75L)+ P_myo-2_::gfp]* was generated and mapped to position I+5.1 (Shin, *et al*., in preparation).

### Fluorescent microscopy

L3 animals were mounted in 5 mM sodium azide/M9 buffer on slides with 3% agar pad. *pmk-1(re170[pmk-1::mNG^3xFlag])* animals were grown on *Comamonas sp*. (DA1877) (Avery and Shtonda, 2003) and mounted in 2 mg/mL tetramisole in M9 buffer on slides with 3% agar pad. We hypothesized that stress caused constitutive translocation to the nucleus. However, growth on *Comamonas sp*. bacteria, which are thought to be non-inflammatory (Avery and Shtonda, 2003), did not alter the degree of nuclear translocation. Similarly, mounting animals on tetramisole rather than sodium azide did not abolish nuclear translocation. All images were captured by A1si Confocal Laser Microscope (Nikon) using NIS Elements Advanced Research, Version 4.40 software (Nikon).

### CRISPR/Cas9-dependent genome editing

Repair templates were generated by PCR amplification from genomic DNA of ~500 bp homology by Q5 polymerase (NEB), digesting of the target SEC vector, and Gibson Assembly (NEB) directed by homologous ends. The sgRNA targeting sequences were inserted into pJW1236, the Cas9+sgRNA (F+E) plasmid (Ward, 2015), by Q5 site-Directed Mutagenesis (NEB). Plasmids, sgRNA sequences including PAM, and repair ssODNs used were listed in Supplementary Tables 3-5. mNG^^^3xFlag::GCK-2 was generated by microinjection of repair template (20 ng/μl), sgRNA-Cas9 #1 (25 ng/μl), sgRNA-Cas9 #2 (25 ng/μl), and injection marker *P_myo-2_::mCherry* (2.5 ng/μl) into wild-type (N2) animals. sgRNA targeting sequences are listed (Table S4). mKate2^3xFlag::RAL-1(+) and mKate2^3xFlag::RAL-1 (G26V) were generated by microinjection of repair template (20 ng/μl), sgRNA-Cas9 #1 (25 ng/μl), sgRNA-Cas9 #2 (25 ng/μl), and injection marker *P_myo-2_::gfp* (10 ng/μl) into N2. The *ral-1(G26V)* mutation was generated by Q5 site-Directed Mutagenesis (NEB) into the homology arm of the repair template used for CRISPR. PMK-1::mNG^3xFlag was generated by microinjection of repair template (10 ng/μl), sgRNA-Cas9 #1 (50 ng/μl), sgRNA-Cas9 #2 (50 ng/μl), and injection marker *P_myo-2_::mCherry* (2.5 ng/μl) into N2. To minimize steric hindrance, the linker sequence N-SAGGSAGGSAGG-C (Komatsu et al., 2011; 5’ -TCAGCCGGAGGTAGCGCCGGCGGAAGTGCTGGTGGA-3) was inserted between *pmk-1* and mNG coding sequences, while RAL-1 and GCK-2 tagging had shorter linker N-SAGG-C (5’ -GGAGCCGGATCT-3’) between FP::epitope and the N-terminus. Animals were handled and treated with 5 mg/ml hygromycin as described (Dickinson et al., 2015). The CRISPR knock-in results were confirmed by genotyping PCR using Taq PCR Master Mix (QIAGEN) and sequencing (Genewiz). Using the previously tagged DV3228 *gck-2(re113[mNG^3xFlag::gck-2])* as a starting point, the *gck-2(mid-Δ)* was generated by Co-CRISPR using *dpy-10(cn64*gf) as a Co-CRISPR marker (Arribere et al., 2014). sgRNA-Cas9 constructs were prepared by Q5 site-Directed Mutagenesis (NEB) of pJW1236. The repair ssODN, providing 35 bases of flaking homology arms on each side of the repaired break, was synthesized by IDT. We microinjected DV3228 with the two sgRNA-Cas9 constructs (each 25 ng/μl), repair oligo (10 μM), sgRNA for *dpy-10(cn64*gf) (pJA58) (25 ng/μl), ssODN repair donor for *dpy-10(cn64*gf) (600 nM), and injection maker *Pmyo-2::mCherry* (2.5 ng/μl). Animals were handled and isolated as described (Arribere et al. 2014) by picking Rols and Dpys for PCR genotyping to detect the deletion, followed by sequence analysis of the repaired region and outcrossing to N2.

We assessed CRISPR tagged alleles for possible impacts on function by crossing tagged alleles into the *let-60(n1046*gf) and the *let-23(sa62*gf) sensitized backgrounds and comparing ectopic 1° induction of *n1046* or *sa62* alone vs. n1046 or sa62+tagged allele strains. We observed induction indices of 1.4 vs. 1.4 for *n1046* vs. *n1046; gck-2(re113)*, respectively (P = 0.9), 1.4 vs. 1.2 for *n1046* vs. *n1046; ral-1(re218)*, respectively (P = 0.3), and 1.3 vs. 1.3 for *sa62* vs. *sa62; pmk-1(re170)*, respectively (P = 0.8). The functional impact of CRISPR putative gain-of-function mutations are shown in Fig. 4B (for the *ral-1(re160*gf) G26V allele) and Fig. 4F (for the *gck-2(re113re222)* mid-Δ allele). By visual inspection, none of the CRISPR tags or mutations altered the wild-type development.

### Western blotting

Animals were lysed in 4% SDS loading buffer by boiling at 90°C for 2 minutes. Protein samples were run on 4-15% SDS gel (BIO-RAD). Monoclonal anti-Flag antibody (Sigma-Aldrich F1804) and monoclonal anti-α-tubulin antibody (Sigma-Aldrich T6199) were diluted 1:2000 in blocking solution. Secondary antibody, goat anti-mouse (MilliporeSigma 12-349), diluted in 1:5000 in blocking solution. ECL reaction (Thermo Fisher Scientific) has done for signal generation. Immunoactive proteins were detected by film processor, SRX-101A (Konica Minolta) on X-ray film (Phenix).

### Bacterially mediated RNA interference

RNAi plasmids used were: III-7M13 *(ral-1)*, V-4P08 *(gck-2)*, X-5G21 *(mig-15), gfp* (Zand et al., 2011), and *luciferase*. For an RNAi negative control, a luciferase fragment not having sequence overlap with the *C. elegans* genome was amplified from SRE-luciferase plasmid and cloned into the HindIII- and XhoI-cut sites of L4440/pPD129.36. The host of bacterially-mediated RNAi clones was HT115 (Timmons and Fire, 1998). RNAi experiments were performed at 23°C on NGM agar plates supplemented with 1 mM IPTG and 50 μg/ml carbenicillin. Plates were seeded with 80 μl dsRNA-producing bacteria, grown overnight at room temperature, then populated with late L4 animals. Parents were transferred to another RNAi plate after 1 day, and ectopic pseudovulvae were scored by DIC at the late L4 stage, 2 days later.

### MiniMos

*reSi6 [P_lin-31_::mKate2::linker::pmk-1::unc-54 3’UTR]* was generated by miniMos (de la Cova et al., 2017; Frokjaer-Jensen et al., 2014). mKate2::linker(SAGG) was tagged to the N-terminal of *pmk-1* with *lin-31* promoter and *unc-54* 3’UTR. MiniMos-based plasmid pSH41 was generated by subcloning of *mKate2::linker::pmk-1* into pCC249 (P*_lin-31_::unc-54 3’UTR)* using Gibson Assembly (NEB). We microinjected N2 wildtype with the *P_myo-2_::gfp* (20 ng/μl), pGH8 (P*_rab-3_:mCherry:unc-54 3’UTR)* (10 ng/μl), pCFJ601 *(P_eft-3_:mos1 transposase::tbb-2 3’UTR)* (65 ng/μl), pMA122 *(P_hsp16.41_::peel-1::tbb-2 3’UTR)* (10 ng/μl), and pHS41 (P*_lin-31_::mKate2::linker::pmk-1::unc-54 3’UTR)* (10 ng/μl). The insertion site is unknown. MiniMos results were tracked by observing mKate2 signal in VPCs.

---

**Figure S1.**
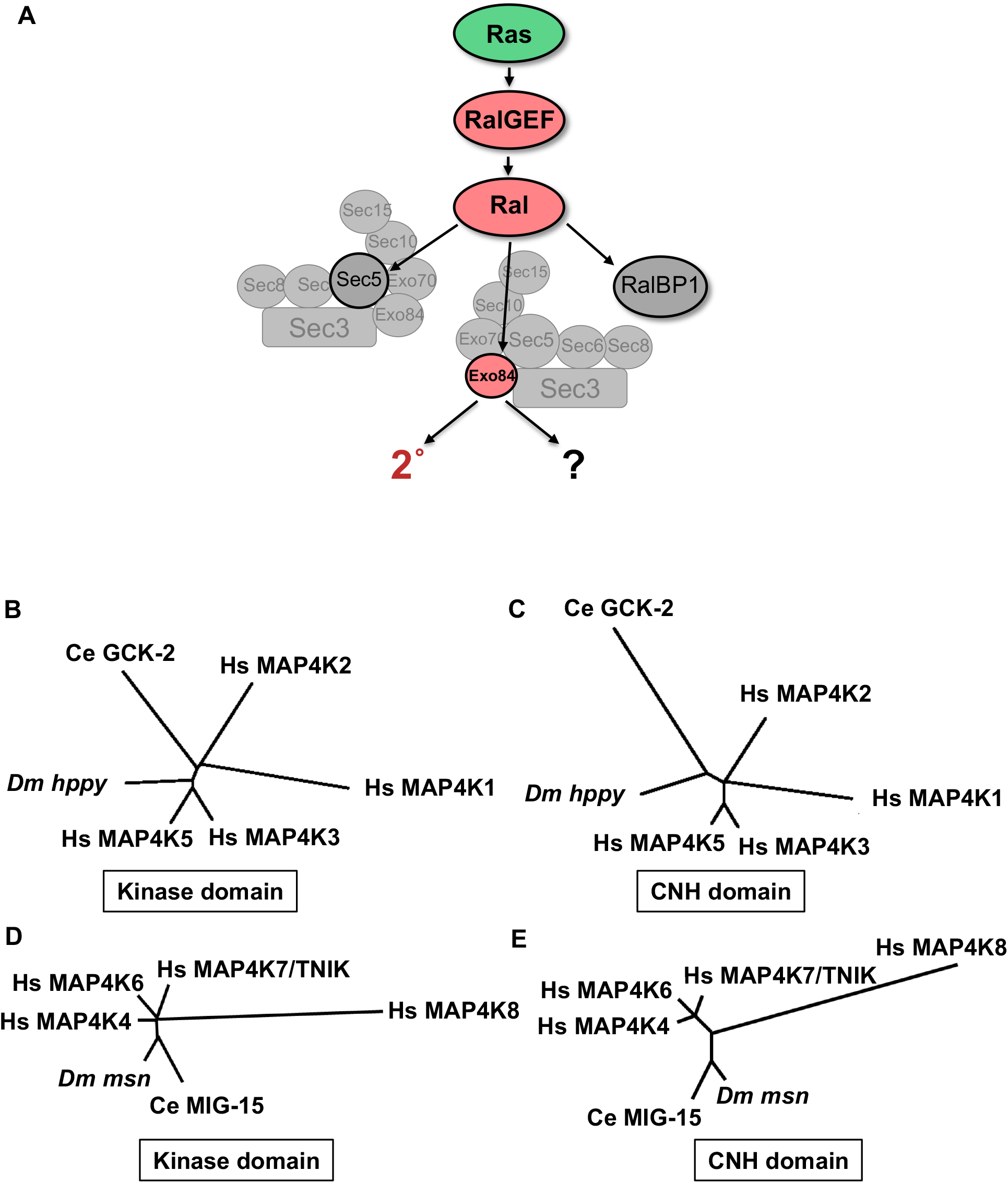
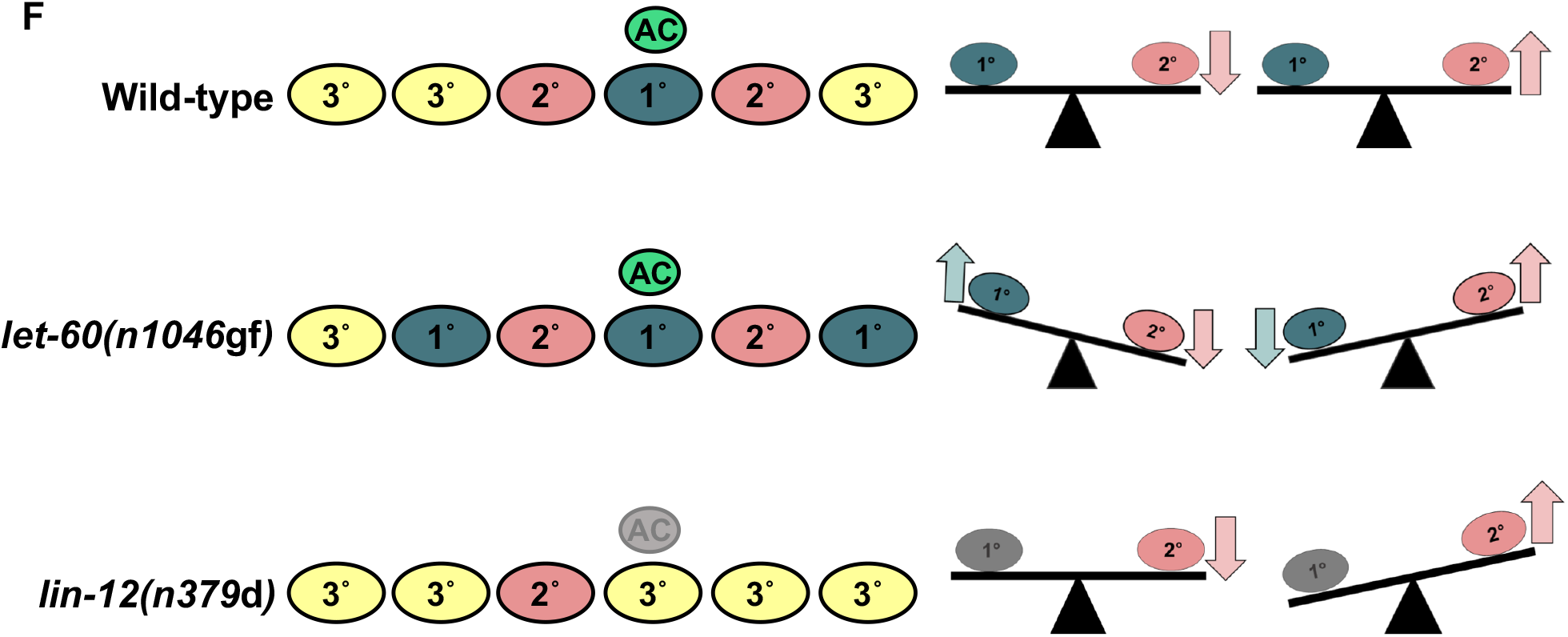
A model of Ral effector use in VPC fate patterning, MAP4K families, and sensitized backgrounds, Related to Figure 1. (A) Ras signals through RalGEF-Ral in cancer and *C. elegans* 2° fate induction. Mammalian Ral has three known oncogenic binding partners: Exo84, Sec5, and RalBP1. (B,C) Unrooted dendrograms of GCK-2/GCK-I subfamily (C. *elegans* GCK-2, *Drosophila happyhour (hppy)*, mammalian MAP4K1/HPK1, MAP4K2/GCK, MAP4K3/GLK, and MAP4K5/GCKR/KHS1) kinase (B) and CNH (C) domains. (D,E) Unrooted dendrograms of MIG-15/GCK-IV subfamily *(C. elegans* MIG-15, *Drosophila misshapen (msn)*, vertebrate MAP4K4/NIK/HGK, MAP4K6/MINK, MAP4K7/TNIK, and MAP4K8/NRK/NESK) kinase (D) or CNH (E) domains, calculated by CLUSTALW. Ce: *C. elegans*, Dm: *D. melanogaster*, and Hs: *Homo sapiens*. Between the GCK-2 and MIG-15 paralogs, the kinase domain and CNH domain share 45% and 29% sequence identity, respectively. Between the GCK-2 and *Drosophila hppy* orthologs, the kinase and CNH domains share 71% and 31% identity, respectively. Between the MIG-15 and *Drosophila msn* orthologs, the kinase and CNH domains share 83% and 79% identity, respectively. Thus, sequence conservation is greater within a subfamily between species than between subfamilies within a species, the definition of subfamily. (F) Schematics of genetically sensitized backgrounds and responses to altered degree of modulatory 2° signaling (after Zand et al., 2011). Wild-type animals have the 3°-3°-2°-1°-2°-3° pattern of vulval cell fates. In the *let-60(n!046*gf) background with ectopic 1° cells, levels of ectopic ?s increase when 2°-promoting signal is blocked and decrease when 2°-promoting signal is activated. In the *lin-12(n379*d) background without an AC (and hence no LIN-3/EGF) and with occasional ectopic 2° cells, levels of ectopic 2° cells are unaltered when 2°-promoting signal is blocked and increased when 2°-promoting signal is activated.

**Figure S2.**
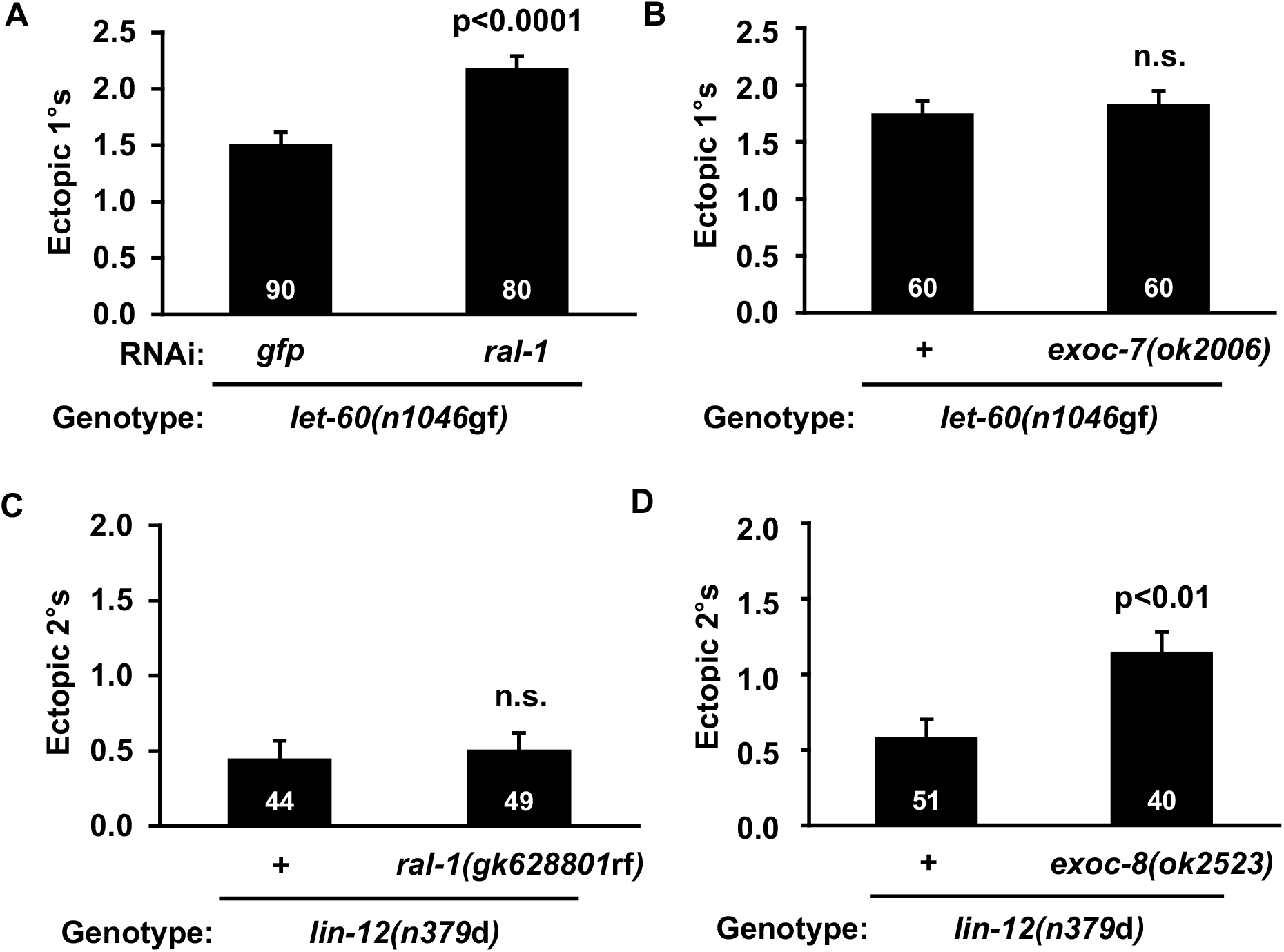
EXOC-8 but not SEC-5 or RLBP-1 functions in VPC fate patterning, Related to Figure 2. Y axes indicate number of ectopic 1° or 2° cells. (A) *ral-1(RNAi)* increased ectopic 1° induction in the *let-60(n1046*gf) background. *gfp*-directed RNAi was used for negative control (Zand et al., 2011). (B) The *exoc-7(ok2006)* deletion did not alter ectopic 1° induction in the *let-60(n1046*gf) background. (C) As predicted, the *ral-1(gk628801)* R139H missense mutation did not alter ectopic 2° induction in the *lin-12(n379*d) background. (D) The *exoc-8(ok2523)* deletion conferred increased ectopic 2° induction *lin-12(n379*d). P values calculated by t test. Error bars = S.E.M.

**Figure S3.**
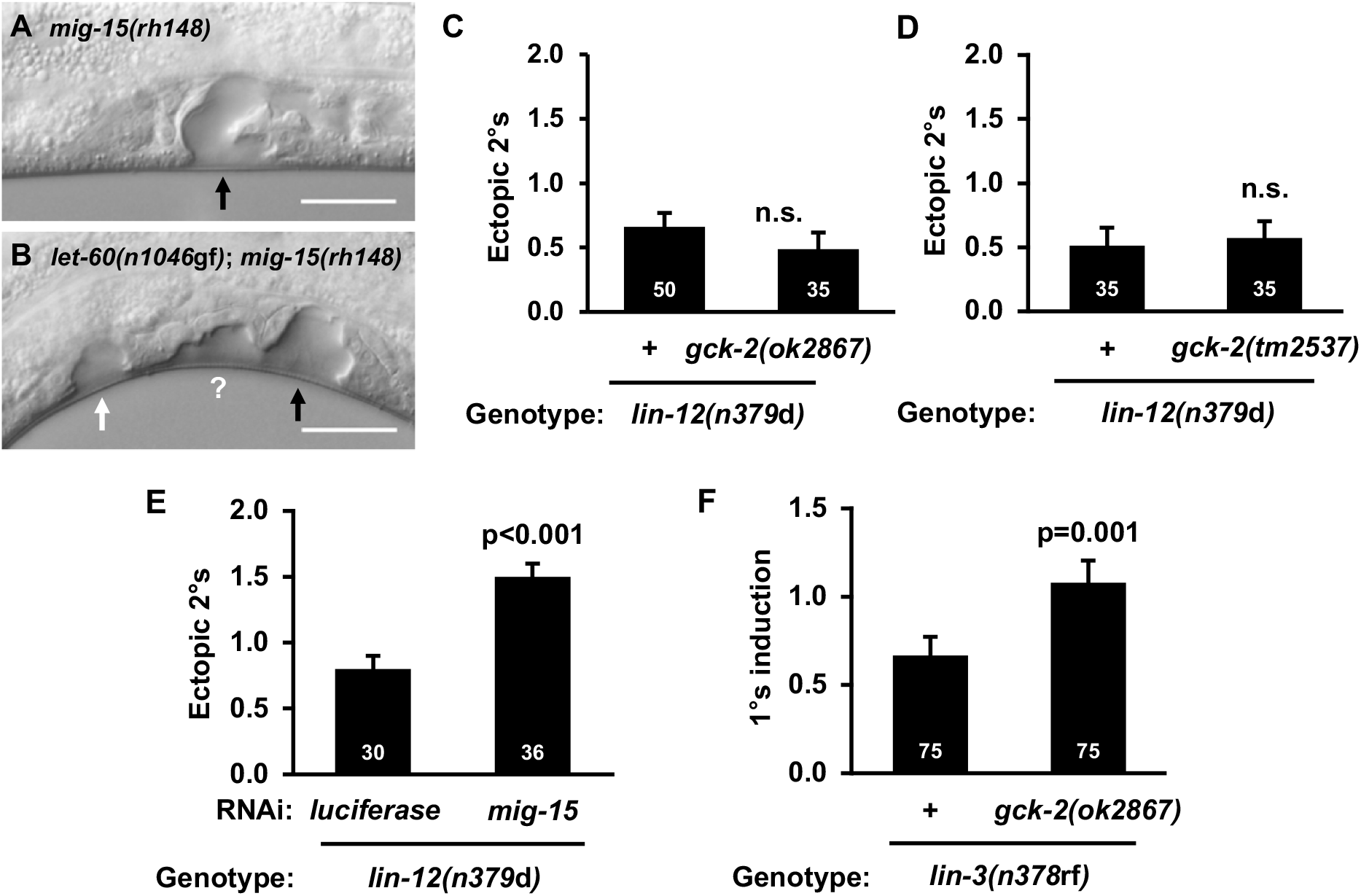
Loss of GCK-2 but not MIG-15 conferred defects consistent with a RAL-1 effector, Related to Figure 3. (A) Representative vulval morphogenetic defects in *mig-15(rh148)* and (B) *let-60(n1046*gf); *mig-15(rh148)*. Vulval morphogenetic defects of *mig-15(rh148)* animals precluded conventional scoring of 1° induction because 2° lineages fail to join the central 1° lineage (see 3G). However, this *mig-15* mutant morphogenetic defect does not confound scoring of isolated 2°s in the *lin-12(n379*d) background, because no migration occurs. Black arrows indicate normal vulvae, white arrows indicate ectopic pseudovulvae, the question mark is probably a 2°, but separated from the main vulva due to morphogenetic defects. (C-D) *gck-2(ok2867)* or *gck-2(tm2537)* does not alter 2° induction (E) while *mig-15(RNAi)* shows increased ectopic 2° induction in the *lin-12(n379*d) background. (F) *gck-2(ok2867)* partially suppresses the absent 1° induction phenotype of *lin-3(n378*rf). P value calculated by *t* test. Error bars = S.E.M.

**Figure S4.**
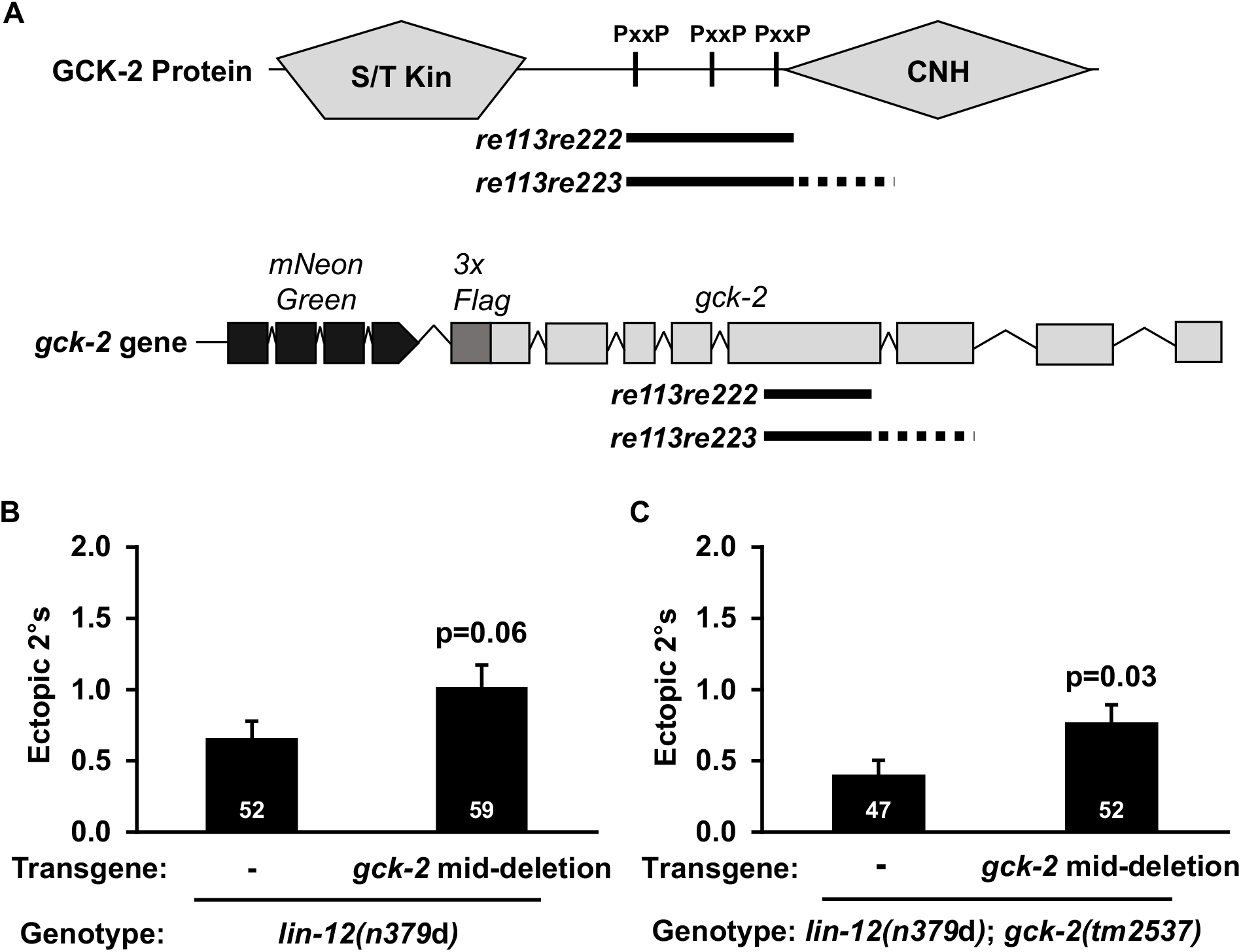
Design and validation of endogenous GCK-2 mid-deletion and GCK-2 mid-deletion transgenics, Related to Figure 4. (A) The *gck-2* mid-deletion removes the central proline-rich region from the gene and protein. The solid black line shows the in-frame deletion while the line that becomes dotted represents the out-of-frame deletion. (B) *reEx143 [P_lin-31_:: gck-2 (mid-deletion)*, *P_myo-2_::gfp]* extrachromosomal array in the *lin-12(n379*d); *gck-2(+)* background. (C) *reEx161[P_lin-31_::gck-2(mid-deletion)*, *P_myo-2_::gfp]* extrachromosomal array in the *lin-12(n379*d);*gck-2(tm2537)* background. P value calculated by *t* test. Error bars = S.E.M.

**Figure S5.**
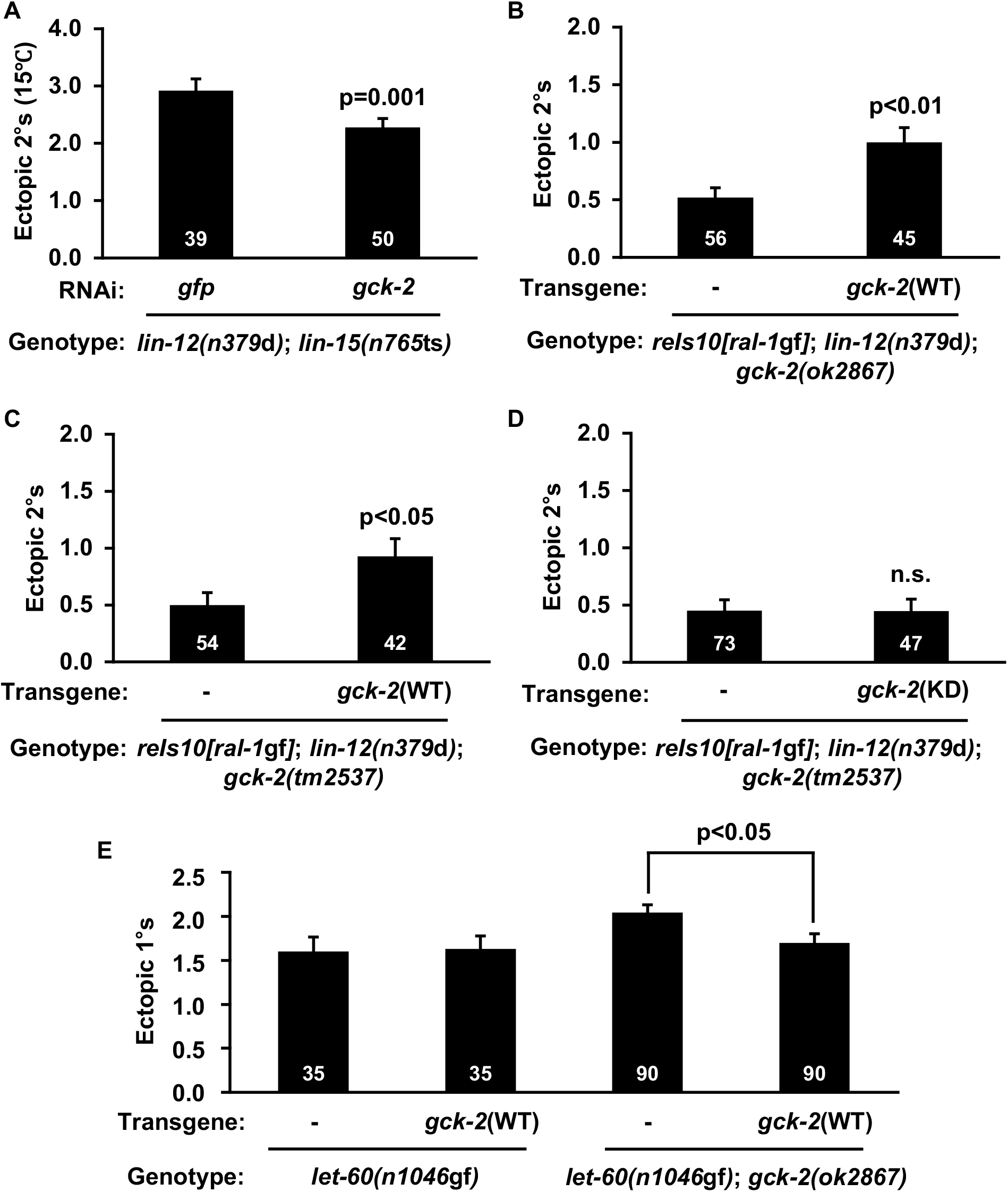
GCK-2 functions cell autonomously downstream of Ral, Related to Figure 5. Strong enhancement of *lin-12(n379*d)-dependent 2° induction by *lin-15(n765*ts) at 15° is inhibited by *gck-2(RNAi)*, with *gfp(RNAi)* as a negative control. (B-D) Additional transgenic arrays to repeat Figs. 5E and 5F (B: *reEx167[P_lin-31_::gck-2(+), P_myo-3_::gfp]*, C: *reEx177[P_lin-31_::gck-2(+), P_myo-3_::gfp]*, and D: *reEx180[P_lin-31_::gck-2(K44E), P_myo-3_::gfp])*. (E) Vulva-specific expression of GCK-2(+) does not alter ectopic 1° induction in the *let-60(n1046*gf) background (first two columns, *let-60(n1046*gf); *reEx113[P_lin-31_:gck-2(+), P_myo-2_::gfp]*), but does rescue the enhancement of ectopic 1° induction by the *gck-2(ok2867)* mutation (second two columns, *let-60(n1046gf); gck-2(ok2867); reEx112[P_lin-31_::gck-2(+), P_myo-2_::gfp]*). Each column pair compares array-bearing vs. nonarray-bearing siblings. P value calculated by *t* test. Error bars = S.E.M.

**Figure S6.**
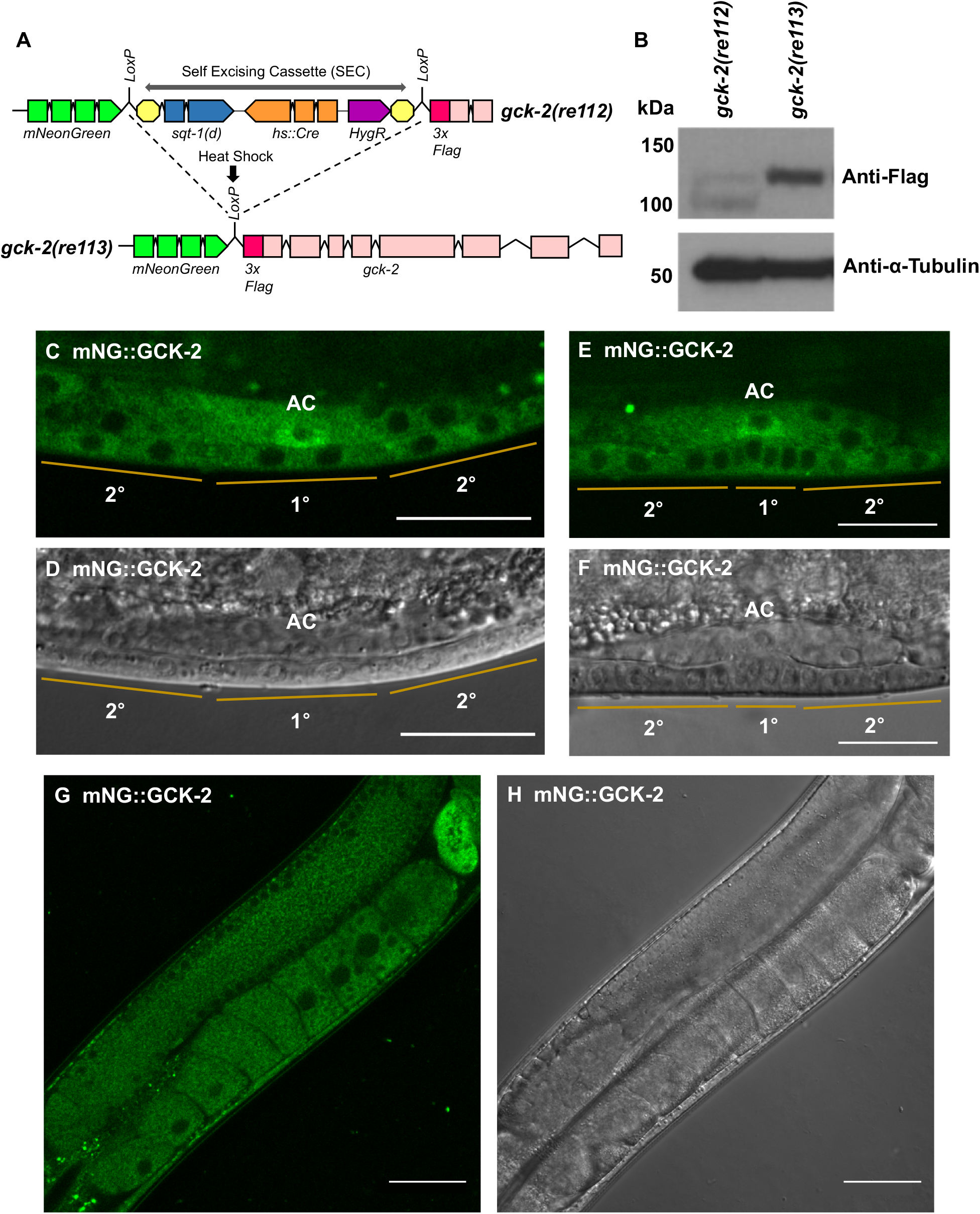
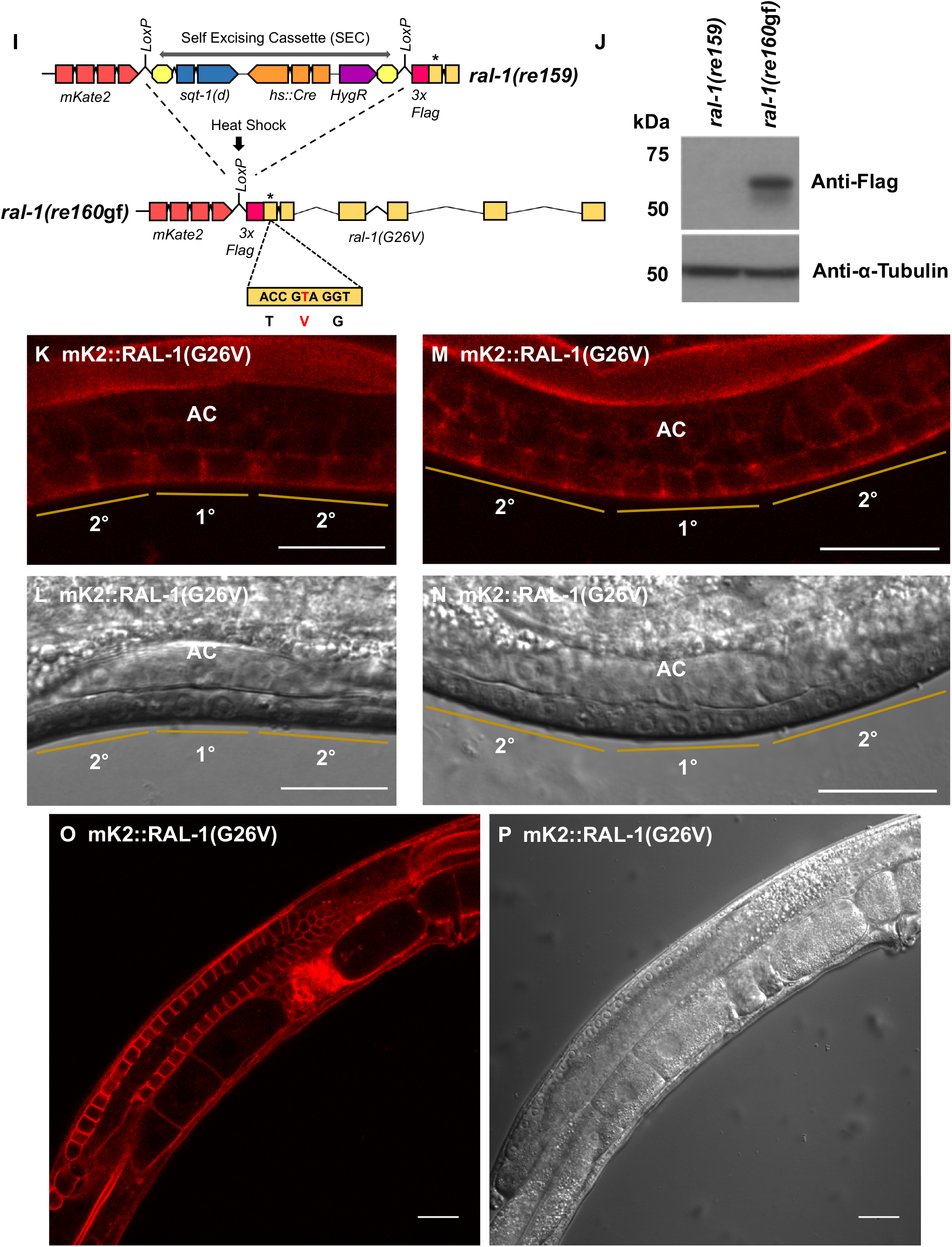
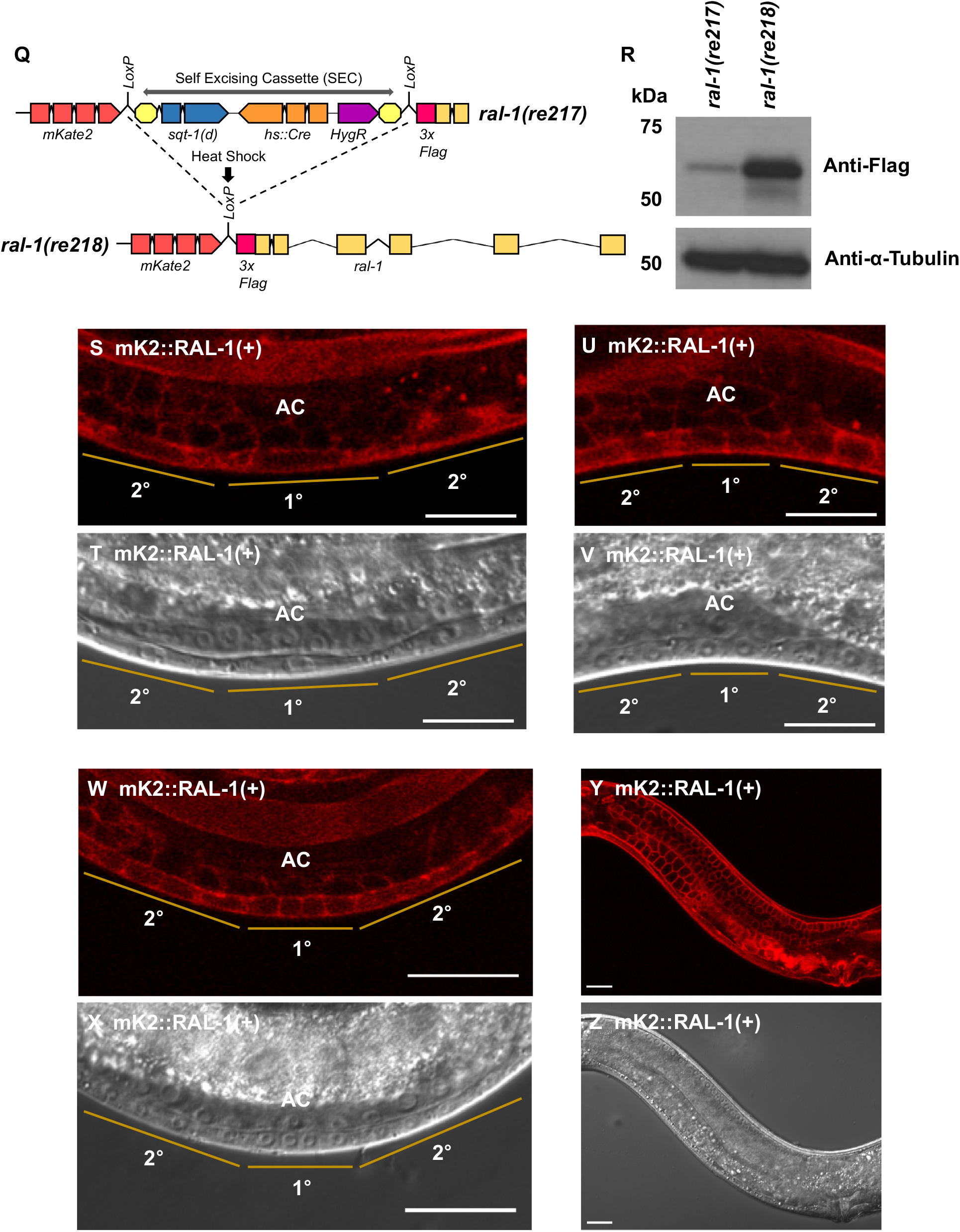
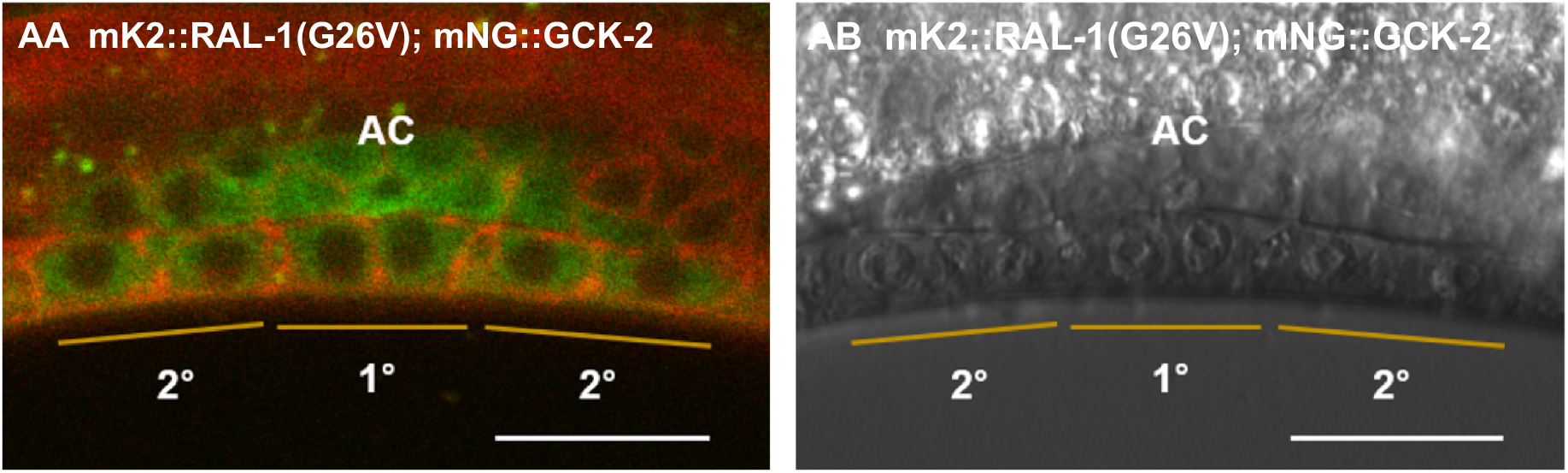
CRISPR knock-in strategies, western blot validations, and localization of endogenously tagged GCK-2, RAL-1(G26V), and RAL-1, Related to Figure 6. (A) The Self-Excising Cassette (SEC) has positive selection marker *HygR* (hygromycin resistance), negative selection marker *sqt-1(*d) (confers a dominant Rol phenotype), and heat-shock Cre to excise the cassette (Dickinson et al., 2015). Before heat shock, animals are selected by Rols that survive hygromycin treatment. After heat-shock, the region between the two LoxP sites is excised, and functional endogenously tagged *gck-2* is generated. (B-H) Endogenously tagged GCK-2. (B) CRISPR knock-in results were confirmed by western blot. First lane: *gck-2(re112[mNG^SEC^3xFlag::gck-2])*, second lane: *gck-2(re113[mNG^3xFlag::gck-2])*. Endogenously tagged GCK-2 protein (mNG::3xFlag::GCK-2) is ~124 kDa, and was detected by anti-Flag antibody (1:2000) (Sigma-Aldrich F1804), with loading control Anti-α-Tubulin (1:2000) (Sigma-Aldrich T6199). “mNG” = mNeonGreen (Shaner et al., 2013). (C-F) Representative confocal and DIC micrographs of the 1° and2° vulval lineages of *gck-2(re113[mNG^3xFlag::gck-2])* animals at the Pn.px (2-cell) and Pn.pxx (4-cell) stages. Tagged GCK-2 is cytosolic in vulval lineages. The Anchor Cell (AC) exhibits strong cytosolic tagged GCK-2. “AC” label is placed directly above the Anchor Cell. (G, H) Representative confocal and DIC micrographs of adult germlines of *gck-2(re113[mNG^3xFlag::gck-2])* animals, exhibiting cytosolic expression. Brighter expression at the right side of the image is the spermatheca. (I) SEC strategy for N-terminal tagging and G26V mutagenesis of endogenous RAL-1 was similar as for N-terminal tagging of GCK-2 (Fig. S6A), except mKate2 was substituted for mNeonGreen and the downstream homology arm sequence contained a missense mutation predicted to confer a G26V change in tagged RAL-1 protein. (J-P) Endogenously tagged RAL-1(G26V). (J) CRISPR knock-in results were confirmed by western blot: First lane: *ral-1(re159[mKate2^SEC^3xFlag::ral-1(G26V)])*, second lane: *ral-1(re160gf[mKate2^3xFlag::ral-1(G26V)])*. Endogenously tagged mKate2::3xFlag::RAL-1(G26V) was detected by anti-Flag antibody (1:2000) (Sigma-Aldrich F1804), showing a protein size of ~56 kDa for RAL-1A, with loading control Anti-α-Tubulin (1:2000) (Sigma-Aldrich T6199). “mK2” = mKate2 (Shcherbo et al. 2009). RNAseq data for *ral-1* in Wormbase predict a potentially longer isoform predicted to run at 61 kD (RAL-1B), but with far less transcript (~13% of total). RAL-1B, if made, would encode an N-terminal extension not found in other species. Since we did not detect an additional band by western blotting, we hypothesize that RAL-1B is simply an extended 5’UTR that is not translated into an alternate isoform. (K-N) Representative confocal and DIC micrographs of the 1° and 2° vulval lineages of *ral-1(re160gf[mKate2^3xFlag::ral-1(G26V)])* animals at the Pn.px (2-cell) and Pn.pxx (4-cell) stages show predominantly localization to plasma membrane and adherens junctions. (O, P) Representative confocal and DIC micrographs of adult germlines of *ral-1(re160gf[mKate2^3xFlag::ral-1(G26V)])* animals, showing plasma membrane localization. The lower left corner of (O) shows the double line of the intestinal lumen, where tagged RAL-1(G26V) is strongly localized to adherens junctions. (Q-Z) Endogenously tagged RAL-1(+). (R) CRISPR knock-in results were confirmed by western blot: First lane: *ral-1(re217[mKate2^SEC^3xFlag::ral-1(+)])*, second lane: *ral-1(re218[mKate2^3xFlag::ral-1(+)])*. Endogenously tagged mKate2::3xFlag::RAL-1(+) was detected by anti-Flag antibody (1:2000) (Sigma-Aldrich F1804), with loading control Anti-α-Tubulin (1:2000) (Sigma-Aldrich T6199). (S-X) Representative confocal and DIC micrographs of the 1° and 2° vulval lineages of *ral-1(re218[mKate2^3xFlag::ral-1(+)])* animals at the Pn.p (1-cell), Pn.px (2-cell), and Pn.pxx (4-cell) stages show predominantly localization to plasma membrane and adherens junctions. (Y, Z) Representative confocal and DIC micrographs of adult germlines of *ral-1(re160gf[mKate2^3xFlag::ral-1(G26V)])* animals, showing plasma membrane localization. (AA-AB) Merged confocal and DIC micrographs of the presumptive 1° (P6.p) and 2° (P5,7.p) lineages of the *ral-1(re160gf[mKate2^3xFlag::ral-1(G26V)]); gck-2(re113[mNG^3xFlag::gck-2])* double mutant at the Pn.px (2-cell) stage. Scale bar throughout = 20 μm.

**Figure S7.**
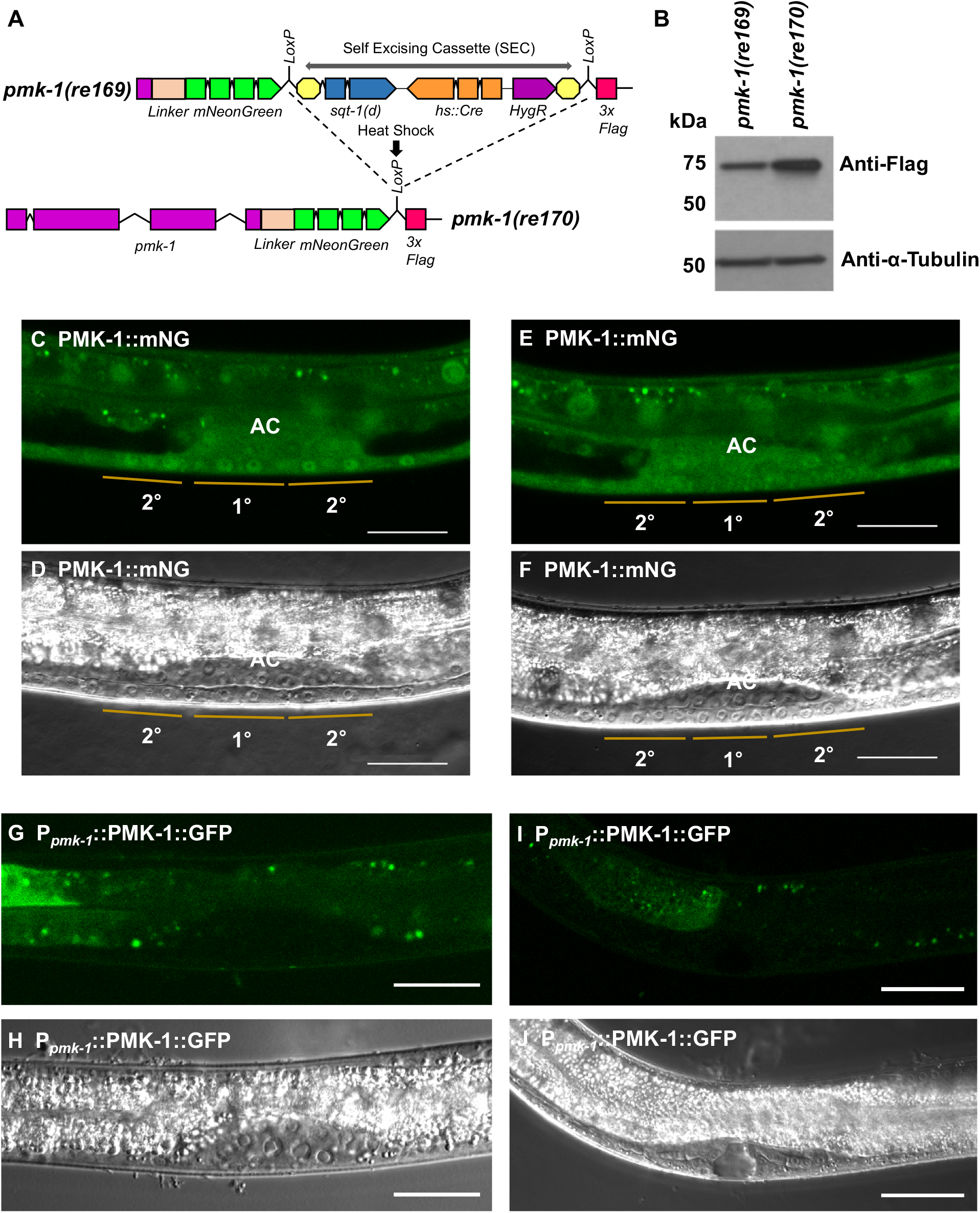
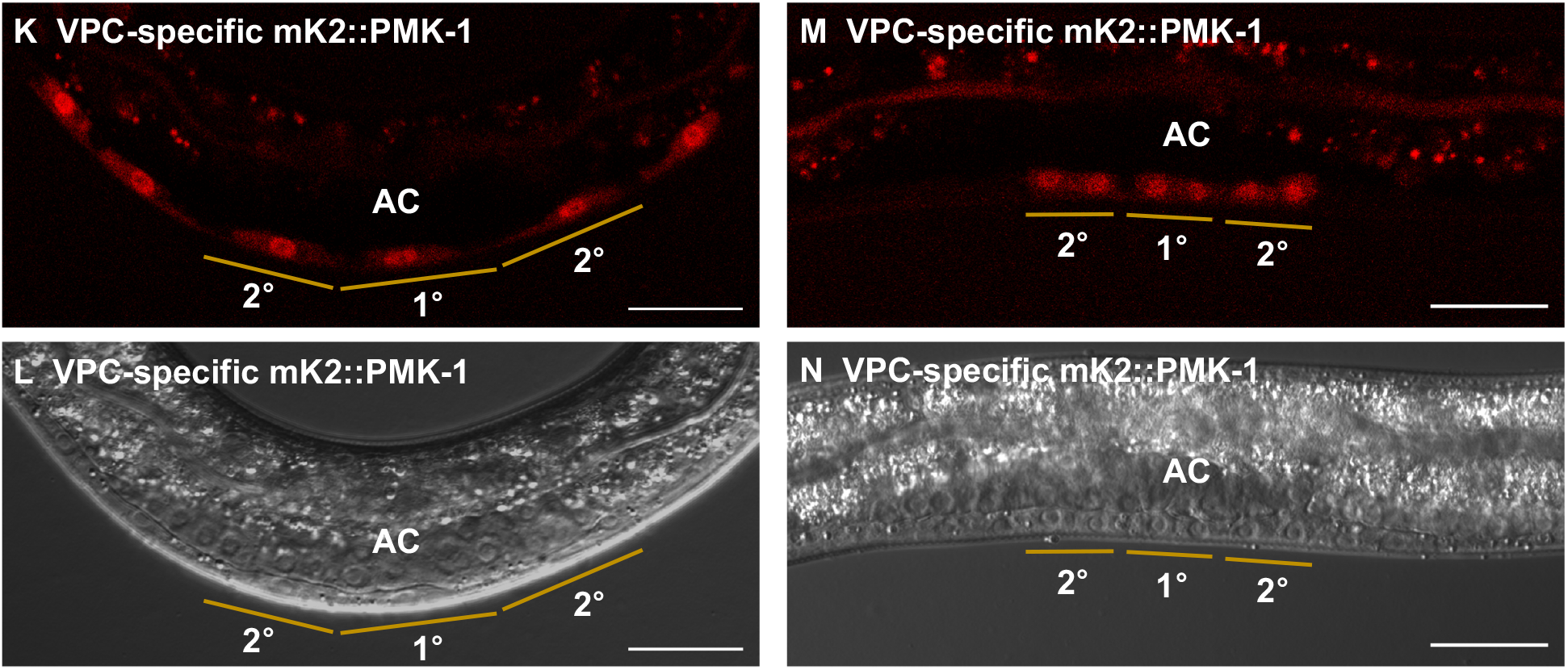
PMK-1 expression and localization, Related to Figure 7. (A) We used a similar CRISPR knock-in strategy for C-terminal tagging of PMK-1 as for N-terminal tagging of GCK-2 and RAL-1 (Figs. S6A, S6I, Q). (B) CRISPR knock-in results were confirmed by western blot: First lane: *pmk-1(re169[pmk-1::mNG^SEC^3xFlag])*, second lane: *pmk-1(re170[pmk-1::mNG^3xFlag])*. Endogenously tagged PMK-1::mNG^^^3xFlag was detected at ~74 kDa by anti-Flag antibody (1:2000) (Sigma-Aldrich F1804), with loading control Anti-α-Tubulin (1:2000) (Sigma-Aldrich T6199). (C-F) Representative confocal and DIC micrographs of the 1° and 2° vulval lineages of *pmk-1(re170[pmk-1::mNG^3xFlag])* animals at the Pn.px (2-cell) and Pn.pxx (4-cell) stages show the expression both in nucleus and in cytoplasm of VPCs. (G-J) confocal and DIC micrographs of P*_pmk-1_::pmk-1 (+)::gfp, rol-6 (su1006)* animals showing expression in intestine, but not in VPCs at L3 stage and vulva at L4 stage. (K-N) Confocal and DIC micrographs of low copy number of *reSi6[P_lin-31_::mKate2::linker::pmk-1::unc-54 3’UTR]* at the Pn.p (1-cell) and Pn.px (2-cell) stages also shows the expression both in nucleus and in cytoplasm of VPCs.

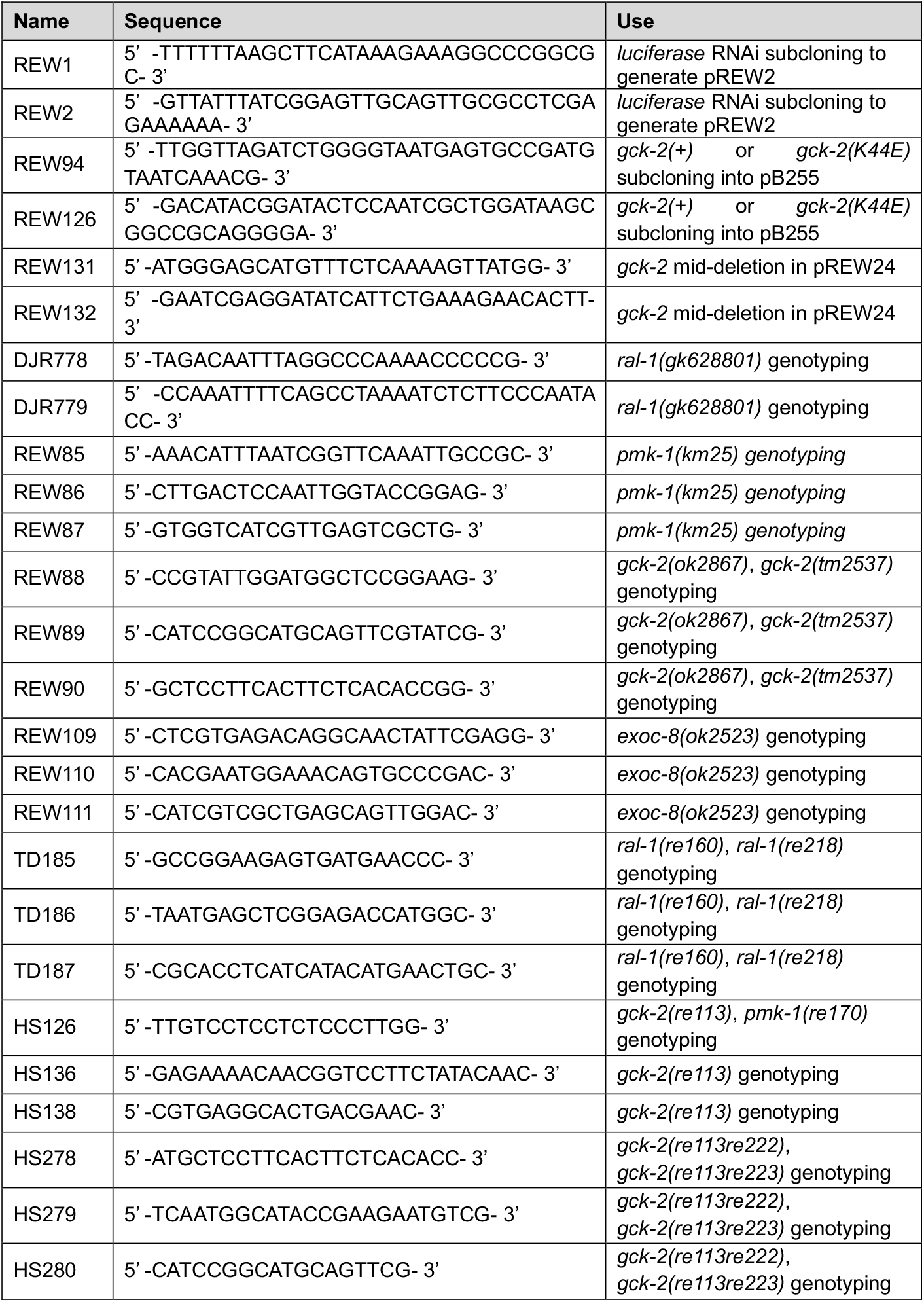
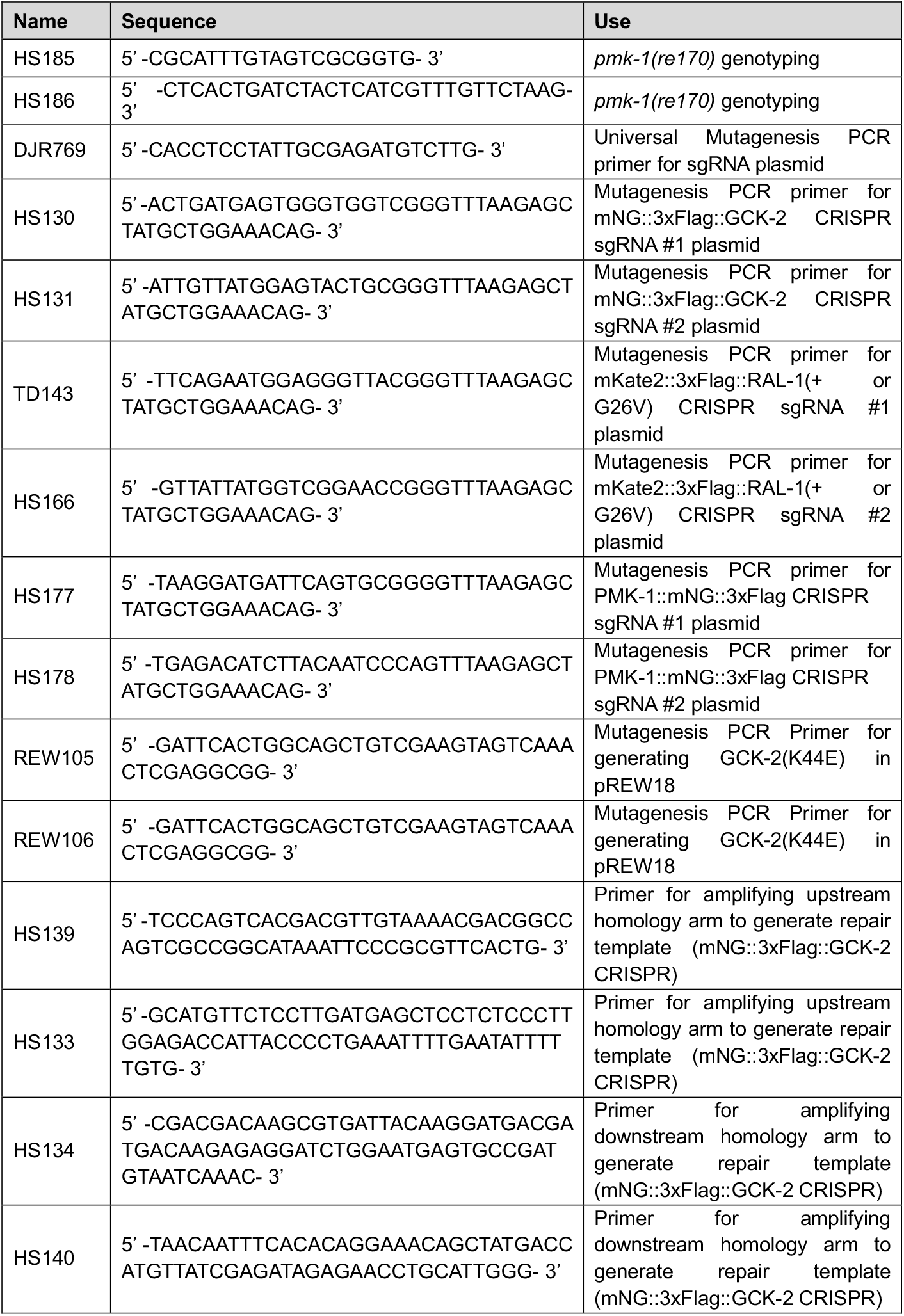
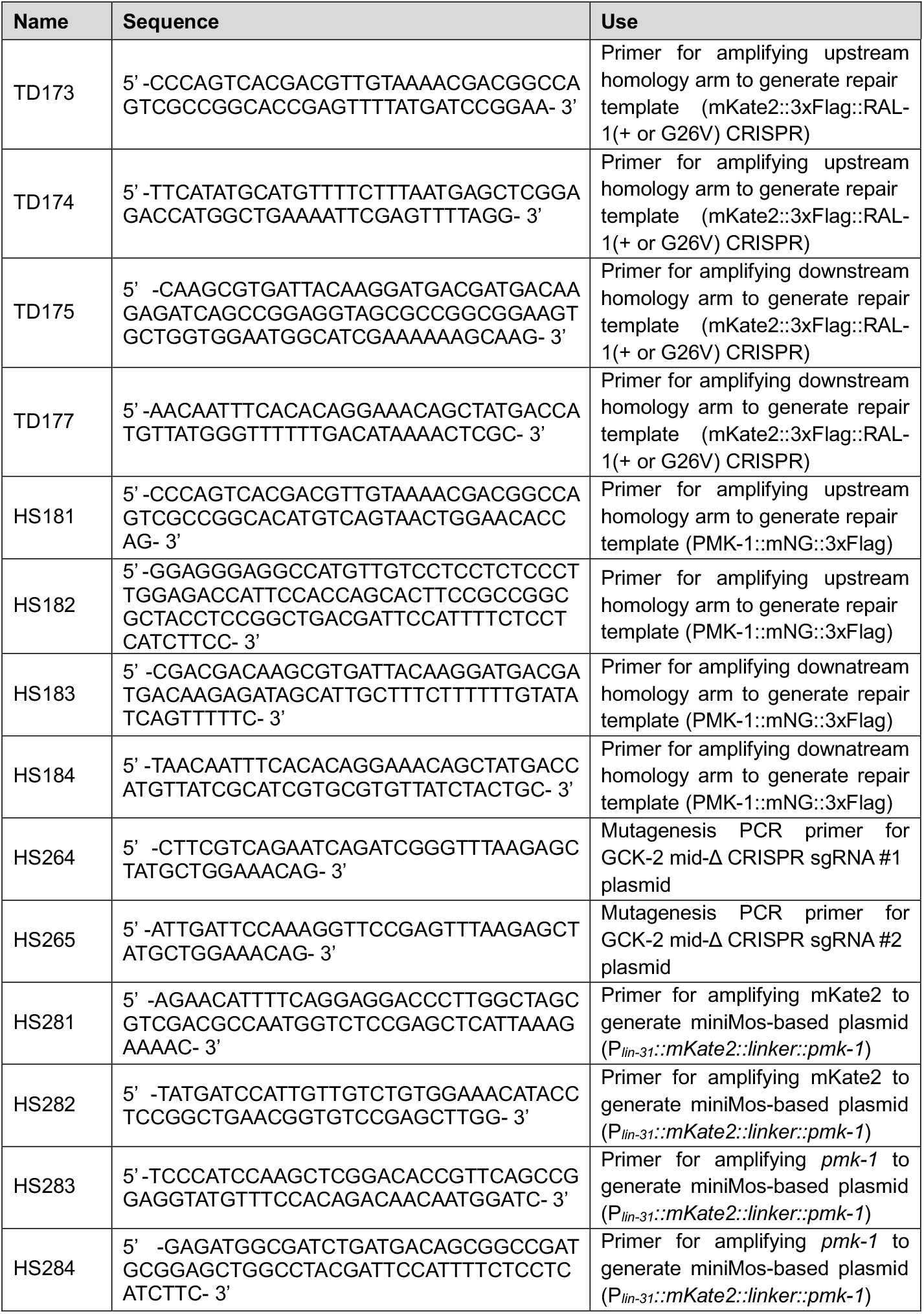
Nucleotides

